# Clinical and molecular features of primary gliosarcoma with digital spatial whole-transcriptome analysis of glial and mesenchymal components

**DOI:** 10.1101/2025.09.02.673845

**Authors:** Matthew D. Wood, Gabriel Zangirolani, Jinho Lee, Tanaya Neff, Kevin Y. Zhang, Christopher L. Corless

**Affiliations:** Department of Pathology and Laboratory Medicine, Oregon Health & Science University, Portland, Oregon, USA; Knight Cancer Institute, Oregon Health & Science University, Portland, Oregon, USA; Johns Hopkins University, Baltimore, Maryland, USA

**Keywords:** Gliosarcoma, glioblastoma, digital spatial profiling, WNT signaling, next-generation DNA sequencing

## Abstract

Gliosarcoma is a rare subtype of IDH-wildtype glioblastoma defined by mixed malignant glial and high-grade sarcomatous histological elements. Gliosarcoma is clinically managed similarly to glioblastoma and has a poor clinical outcome. The sarcoma-like regions of gliosarcoma are thought to represent extreme mesenchymal metaplasia of neoplastic glial cells. Factors contributing to this phenomenon are not completely understood. Here we report a single-institution series of 37 gliosarcomas including next-generation sequencing data on 25 cases and digital spatial whole-transcriptome analysis on 4 cases to characterize differential gene expression between glial and mesenchymal components. Gliosarcoma demographic and genetic features were compared to a cohort of 75 primary adult hemispheric IDH-wildtype non-sarcomatous glioblastomas. Patient age, tumor location, sex, and overall survival in gliosarcoma were similar to glioblastoma. Gliosarcomas showed a significantly lower rate of *EGFR* amplification and a higher rate of *NF1* mutation compared to glioblastomas in next-generation sequencing analysis. Digital spatial whole-transcriptome analysis showed a distinct transcriptomic profile in sarcomatous regions with over-expression of genes involved in extracellular matrix development and remodeling. Selected differentially expressed transcripts were examined further by immunohistochemistry. The glial elements of gliosarcomas showed higher immunoreactivity for Chitinase-3-like protein 1 (CHI3L1) than glioblastomas, but low to absent expression within the sarcomatous elements. Lymphoid Enhancer-Binding Factor 1 (LEF1) immunoreactivity was identified within sarcomatous regions of gliosarcoma without detectable nuclear β-catenin, suggesting a role for β-catenin independent wingless (WNT) effector signaling in sarcomatous transformation. This study adds to the growing literature demonstrating differences in the genetic underpinning of gliosarcoma and glioblastoma, establishes feasibility of spatial transcriptomic approaches in gliosarcoma, and validates digital spatial profiling-based results as a discovery platform to identify pathways and immunohistochemical markers for further study.

## 1 Introduction

Gliosarcoma (GSC) is defined as a subtype of glioblastoma, IDH-wildtype, CNS WHO grade 4 (GBM) in the current World Health Organization Classification of Central Nervous System (CNS) Tumors (41). GSC accounts for about 2% of GBM cases and represents 0.5% of brain tumors overall (18). Histologically, GSC is defined by a pattern of interdigitating malignant glial and high-grade sarcomatous elements, the former regions showing immunoreactivity for glial markers and the latter regions elaborating a reticulin-rich extracellular matrix with low glial marker immunoreactivity (41). This pattern may be seen at initial tumor sampling (primary GSC) or in recurrent tumors after treatment (secondary GSC). Following a seminal description of 3 cases published in 1955, the sarcomatous regions of GSC were presumed to arise from malignant transformation of microvascular proliferation, thus representing a collision event between a sarcoma developing within -- and secondary to -- GBM (12). In the mid-1990’s molecular studies on microdissected regions of glial and sarcomatous regions of GSC showed shared gene mutations and chromosome copy number alterations, supporting a clonal neoplastic process and leading to the current conception that GSC represents a phenotypic change in GBM with tumor cells undergoing extreme mesenchymal metaplasia (5, 6). The distinctive histological pattern of GSC has not been translated to clinically impactful information; treatment approaches for GSC do not differ from modalities used in GBM, and most studies have shown similar overall survival. There are reports of extra-cranial metastasis from GSC (3, 15). This suggests distinct biological behavior, but it remains a rare phenomenon.

Molecular testing is a routine part of clinical neuro-oncologic neuropathology practice, and several studies have reported that the genetic features of GSC overlap with -- but bear some difference from -- typical GBM (41). Studies have differed in their precise findings, but in general have consistently shown a lower rate of *EGFR* alterations in GSC compared to GBM, with some studies also showing higher rates of *NF1*, *TP53*, *PTEN*, and *RB1* alterations in GSC (9, 23). Histologic studies have established that sarcomatous regions of GSC have increased expression of epithelial-mesenchymal transition (EMT)-related transcription factors, matric metalloproteinases, and in a subset of cases nuclear β-catenin localization (27, 28, 37). Given the dramatic biphasic nature of GSC, it is an interesting subtype to study for candidate tissue markers that may give insight into the sarcomatous phenotypic transformation. Such findings could translate to other CNS tumors with sarcomatous features such as IDH-mutant oligosarcoma, sarcomatous IDH-mutant astrocytoma, ependymosarcoma, and CNS tumors with *PATZ1* fusion (2, 19, 35, 39).

Advanced methods for spatial transcriptomic and proteomic analysis of formalin-fixed paraffin embedded (FFPE) tissue can give insight into spatially heterogeneous tumors (16, 24). Singe-cell molecular testing has added an abundance of data on GBM cellular heterogeneity and the impact of non-neoplastic inflammatory and other microenvironment cells on tumor behavior and evolution under treatment (31, 38). In 2008 The Cancer Genome Atlas study identified 4 glioblastoma subtypes based on gene expression profiling of bulk tumor tissue, including a “mesenchymal” subtype enriched for *EGFR*-wildtype, *NF1*-mutated or -deleted cases (7). Subsequent work showed that individual GBM tumors contain a mixture of tumor cells representing various transcriptomically-defined states, and that the cellular state is dynamic and subject to evolution under treatment and/or in response to microenvironmental factors (33). The GBM cellular state provides a powerful framework for evaluating gene expression patterns and determining how histologically distinct regions of these tumors relate to underlying tumor biology. In this study we assembled an institutional cohort of GSC cases, collected their demographic, survival, and genetic features for comparison to GBM, and performed digital spatial whole-transcriptome analysis to identify protein markers evaluable in FFPE tissue sections to gain insight into the biological processes associated with sarcomatous transformation.

## 2 Materials and Methods

### 2.1 Cohort Selection

With Internal Review Board approval, pathology records at Oregon Health & Science University (OHSU) were searched for the term “gliosarcoma” appearing in diagnostic reports from 1/1/1990 to 12/31/2023. Reports for 88 cases from 66 patients were identified and reviewed. Residual/recurrent tumors and cases with ineligible final pathologic diagnoses were excluded. The study cohort of 37 primary gliosarcomas included 24 in-house and 13 consultation cases. Thirty-two (32) cases had a definitive diagnosis of GSC and 5 were designated as possible, consistent, or compatible with GSC. A reference cohort of 75 glioblastomas was assembled from the investigator’s records on in-house and consultation cases of primary adult diffuse gliomas diagnosed in 2021-22, which were profiled by next-generation DNA sequencing as routine practice. Demographic information and clinical outcomes were extracted from the electronic medical record. Overall survival was calculated from the date of the diagnostic neurosurgical procedure to death or censored at the last documented clinical follow-up.

### 2.2 Histology and Immunohistochemistry

Diagnostic slides from GSC cases were reviewed by a board-certified neuropathologist (MDW) to confirm the diagnosis, based on the hematoxylin and eosin (H&E) findings of a high-grade astrocytic glioma with prominent mesenchymal features showing at least focal interdigitation of malignant glial/astrocytic and mesenchymal/sarcomatous elements. Stains for glial fibrillary acidic protein (GFAP), reticulin, and other markers were reviewed when available.

Immunohistochemistry for LEF1 was performed with clone EP310 (Cell Marque, predilute) on a Ventana Ultra stainer with DAB Ultra View detection kit. Immunohistochemistry for CHI3L1 was performed using CHI3L1 antibody (Proteintech 12036-1-AP), antigen retrieval with citrate buffer pH 6.0 at 80° C for 45 minutes, blocking by 3% milk solution for 15 minutes, and primary antibody 1:500 overnight at 4° C. Secondary antibody (Vector Laboratories BA-1000 1:800) was applied for 1 hour at room temperature and detected by incubation with DAB solution.

### 2.3 Genetic and Transcriptomic Analysis

Next-generation DNA and RNA sequencing (NGS) of 10 GSC and 75 GBM samples was performed using amplicon-based panels detailed previously, covering 225 genes for sequence alterations and 21 genes for partner-agnostic fusion detection (44). During this study our molecular panel was expanded to cover 281 genes for sequencing and 43 genes for fusion. The expanded panel was applied to another 15 gliosarcomas and genes with shared coverage on both panels were evaluated. Sequence alterations were interpreted using standard guidelines for variant interpretation in cancer (22). Chromosome copy number alterations were evaluated by visual inspection of genome-wide copy number plots extrapolated from NGS results. Amplification of target genes/regions was defined by a tumor-corrected copy number of >5 copies. Homozygous deletion was defined by a tumor-corrected copy number of <0.8 copies. *MGMT* promoter methylation testing was performed in a CAP/CLIA certified laboratory using methylation-specific PCR after bisulfite conversion. NGS and MGMT tests were done on tumor-rich macrodissected material from unstained slides.

Digital spatial profiling (DSP) analysis was performed according to Nanostring, Inc. technical procedures. In summary, freshly cut 5 μm unstained sections from FFPE tissue were deparaffinized and incubated with Nanostring’s whole transcriptome atlas probes at 37°C for 16 hrs. After stringent washes, tissues were stained with GFAP (AF647, Novus Cat# NBP1-05197), vimentin (AF594, Santa Cruz, Cat# sc-373717), and Syto-13 (Thermo Fisher Scientific Cat# S7575) at room temperature for 1 hour. Regions of interest (ROIs) were selected by two of the investigators (MDW and JL) from at least 3 spatially disparate regions per case. Viable tumor was determined by nuclear Syto-13 staining, glial/astrocytic regions were identified by GFAP fluorescence signal, and sarcomatous regions were identified by absence of GFAP fluorescence signal combined with vimentin signal, elongated nuclear morphology, and fascicular/spindled architecture. UV-cleaved oligonucleotides from each ROI were collected in a 96-well plate and NGS libraries were prepared for sequencing. After sample sequencing, the read data were imported into Nanostring DSP software and processed with quality control, gene filtering, and Q3 normalization for analysis.

### 2.4 Database and Literature Data Retrieval

The Cancer Genome Atlas (TCGA) data were accessed through cBioPortal with the following parameters: dataset - Glioblastoma Multiforme (TCGA, Firehose Legacy), genomic profiles - Mutations and Putative Copy Number Alterations (N=273), search genes *EGFR*, *NF1*, *TP53*, *RB1*, *PTEN*, *PIK3CA*, *PIK3R1*, and *PTPN11*. Thirteen isocitrate dehydrogenase (IDH)-mutant tumors were removed due to their reclassification as astrocytoma, IDH-mutant, CNS WHO grade 4 by current diagnostic criteria (40). Review of the diagnostic reports and/or scanned histology slides identified 10 gliosarcomas, leaving 250 IDH-wildtype non-sarcomatous GBM samples. Genetic data from Chen *et al.* was extrapolated from figure 1(A) (9). Genetic data from Lucas *et al* was taken from annotated data generously provided by the study author.

**Figure 1:**
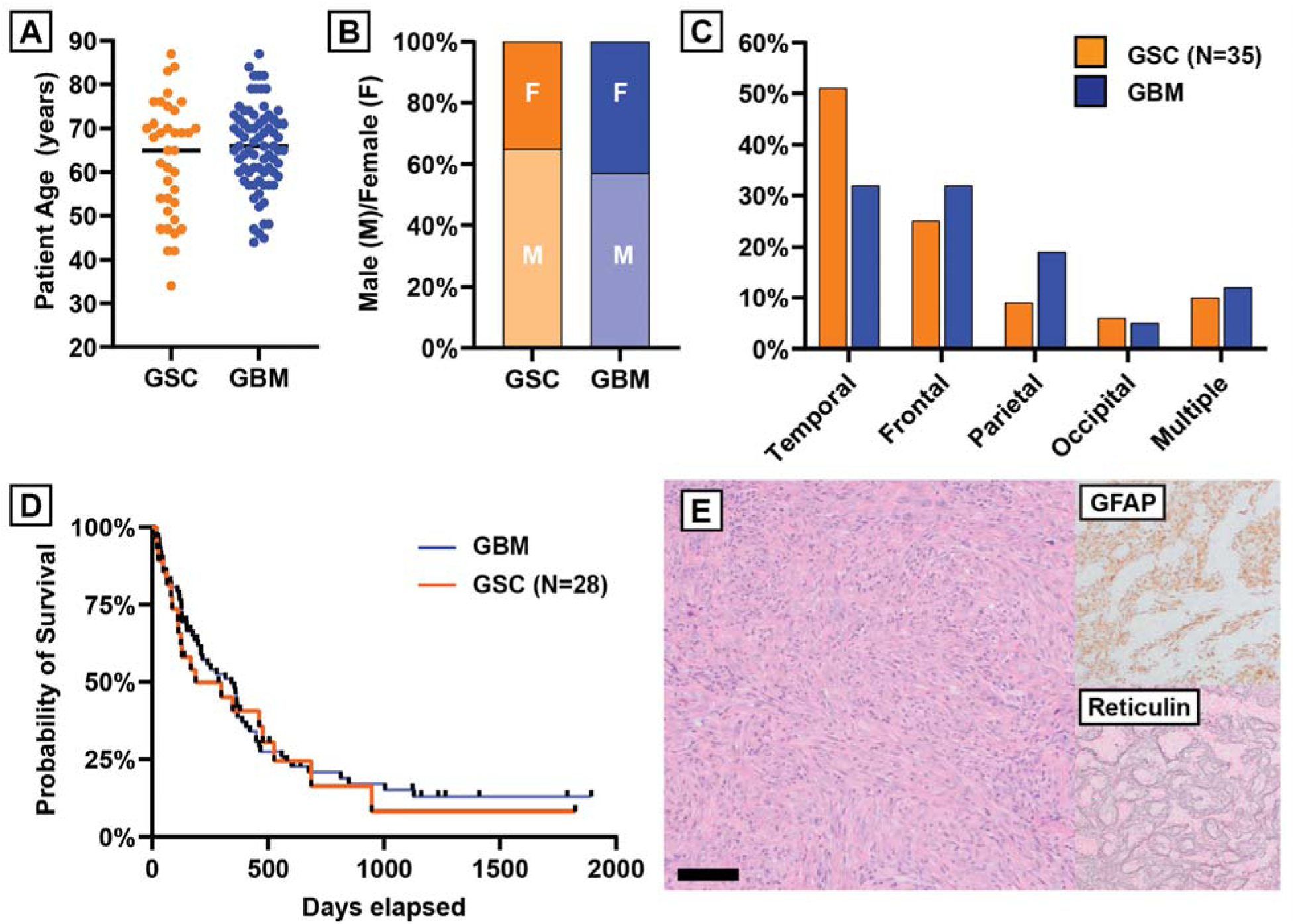
Demographics, overall survival, and representative histology in gliosarcoma and glioblastoma cohort. (A) patient age at diagnosis, (B) patient sex, (C) tumor location, and (D) overall survival. Glioblastoma data is from 75 cases and gliosarcoma data is from 37 cases, unless noted otherwise. (E) Representative gliosarcoma histology (scale bar 100 μm).

### 2.5 Statistics

Statistical analysis was performed using GraphPad Prism 10. Categorical demographic variables were compared using chi-square tests. Age was compared using Welch’s t test. Overall survival was compared using log-rank (Mantel-Cox) test. Gene alteration frequencies from NGS were assessed by 2-tailed Fisher’s exact test with Benjamini-Hochberg correction at false discovery rate 0.05. Gene set enrichment analysis was performed using WEB-based GEne SeT AnaLysis Toolkit (WebGestalt) and the Reactome functional database (11, 46). GeoMx DSP whole-transcriptome analysis (WTA) data were analyzed in R software. Differentially Expressed Genes (DEGs) between sarcomatous and astrocytic regions were identified using the Limma package. Principal Component Analysis (PCA) was performed using the StandR package.

## 3 Results

### 3.1 Demographic, survival, and genetic features in an institutional cohort of primary adult gliosarcoma

The GSC cohort included 24 male and 13 female patients with median age of 66 years (range 34 to 87 years). Tumor location was available for 35 cases with the temporal lobe being most common (N=18; 51%) followed by frontal (N=9; 26%), parietal (N=3; 9%), and occipital (N=2; 6%) lobes. Three tumors involved multiple lobes, with one each of frontal/parietal, frontal/temporal, and temporal/parietal involvement. The GBM cohort included 43 male and 32 female patients with median age 66 years (range 44 to 87 years) with tumors located equally in the frontal or temporal lobes (N=24 each; 32%) followed by the parietal (N=14; 19%) or occipital (N=4; 5%) lobes and 9 involving multiple lobes (12%). Patient age, sex, and tumor location did not statistically differ between GSC and GBM (**Figure 1A-1C**). Clinical follow-up was available for 28 GSC and 75 GBM patients. Median overall survival was 188 days in GSC and 343 days in GBM, and did not reach statistical significance (**Figure 1D**, hazard ratio 1.109, 95% confidence interval 0.658 to 1.870, p-value 0.689). The dominant mesenchymal elements of most gliosarcoma cases corresponded to a high-grade spindle cell sarcoma (68%, **Figure 1E**) with a subset of cases showing lower cellularity malignant spindled cells on a myxoid background (19%) and rare cases showing other patterns such as chondroid differentiation or features resembling exuberant microvascular proliferation (13%).

Genetic features of 25 GSC and 75 GBM cases were compared using an NGS platform detecting sequence alterations, gene fusions (partner agnostic) and splice variants, and copy number alterations. Detailed NGS findings in the GSC cohort are provided as **Supplemental Table S1**. None of the sequenced gliosarcomas had detectable *IDH1* or *IDH2* mutations. The most frequent sequence alterations in GSC were *TERT* promoter region mutation (92%), *PTEN* mutation or homozygous deletion (64%), *TP53* mutation (56%), and *NF1* mutation (48%) (**Figure 2A**). Only 3 gliosarcomas (12%) had *EGFR* amplification, vIII splice alteration, and/or activating mutation. Identification of *BRAF* p.V600E mutation, *CDKN2A* homozygous deletion, and *TERT* promoter region mutation in GSC-25 raised consideration of an (epithelioid) pleomorphic xanthoastrocytoma, CNS WHO grade 3. However, the additional presence of *PTEN* mutation and combined trisomy 7 and monosomy 10, the histologic features including regions of infiltrative growth and absence of eosinophilic granular bodies, and an aggressive clinical course with overall survival of <3 months supported classification of this case as GSC and is consistent with prior reports of GSC sometimes harboring *BRAF* mutation (45).

**Figure 2:**
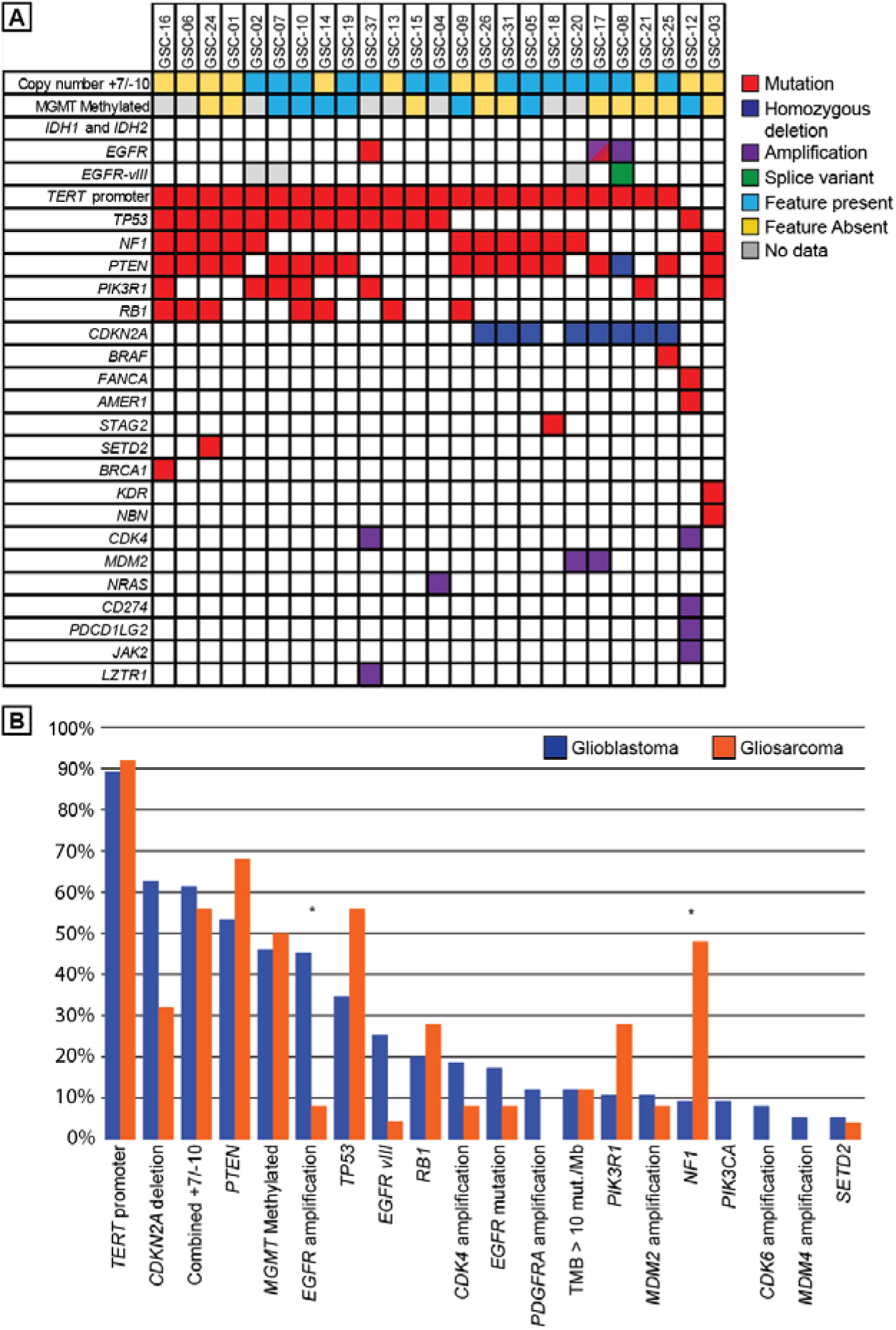
Genetic features of gliosarcoma compared to glioblastoma. (A) OncoPrint summary of genetic features in 25 gliosarcoma cases. (B) Comparison of genetic features of glioblastoma (blue bars, N=75) to gliosarcoma (orange bars, N=25); gene names with unspecified alteration type represents mutations and deletions. Alterations are ordered by frequency in the glioblastoma cohort. * = statistical significance at p<0.05 (see methods).

Comparison of GSC genetic features to the GBM cohort revealed a significantly higher frequency of *NF1* alterations (48% vs. 9%) and a significantly lower frequency of *EGFR* amplification (8% vs. 45%) in GSC (**Figure 2B**). Differences in the frequency of *CDKN2A* homozygous deletion (32% in GSC vs. 63% in GBM; p=0.0105) and in *EGFR*-vIII splice variant (4% in GSC vs. 25% in GBM; p=0.0374) were not significant after correction for multiple hypothesis testing. There were non-significant trends toward higher frequencies of *TP53* (56% vs. 35%; p=0.0977) and *PIK3R1* (28% vs. 11%; p=0.0514) alterations in GSC. Overall, the genetic features in our cohort are consistent with other recent studies on primary and secondary GSC (9, 23).

### 3.2 Gliosarcoma is enriched for *EGFR*-unaltered tumors with *NF1*, *TP53*, and PI3 kinase pathway alterations

To determine if a certain constellation of molecular features is associated with GSC, we combined our dataset with two recent studies including NGS and compared the frequencies of different combinations of alterations to GBM cases from the same studies (when available) and TCGA data (**Table1**). In total, this analysis used the genetic features of 90 gliosarcomas including 63 primary gliosarcomas (25 from this report, 29 from Chen *et al.*, and 9 from TCGA), 7 secondary gliosarcomas (Chen *et al.*), 17 glioblastomas that progressed to GSC at recurrence (Lucas *et al*), and 3 gliosarcomas of unknown primary/secondary status (2 from Chen *et al.* and 1 from TCGA). The comparison group was 250 non-sarcomatous IDH-wildtype GBM cases from TCGA, 75 primary glioblastomas (this study), and 84 glioblastomas that did not transform to GSC at recurrence/progression (Lucas *et al.*). The frequency of an *NF1*-altered, *EGFR*-unaltered tumor was 44.0% in the GSC group (40 of 90), versus 9.0% in the GBM group (37 of 409 - chi-square 70.83, p<0.00001). Occurrence of an additional *TP53* mutation was rare in GBM, identified in 10 of 409 cases (2.4%) compared to 18 of 90 cases of GSC (20.0% - chi-square 42.923; p<0.0001). The constellation of *NF1*, *TP53*, and a PI3K pathway alteration - operationally defined as the presence of *PIK3R1*, *PIK3CA*, or *PTPN11* mutation and/or *PTEN* mutation/deletion (Chen *et al.* incorporating only *PTEN* mutation/deletion) - in an *EGFR*-unaltered tumor occurred in 14 of 90 gliosarcomas (15.5%) versus 4 glioblastomas (1.2% - chi-square 41.3761, p<0.00001). The sensitivity and specificity of this constellation for GSC was 15.5% and 98.7% respectively, with positive predictive value 73.6% and negative predictive value 84.1%. While this is not sufficiently sensitive or predictive to be useful in diagnostic practice, this analysis suggests that a particular molecular background may predispose to GSC developing either at presentation or in the post-treatment recurrence setting.

### 3.3 Digital spatial transcriptomic analysis identified distinct transcriptomic features in malignant glial versus sarcomatous regions of gliosarcoma

Four GSC samples were examined for transcriptomic differences between the gliomatous and sarcomatous regions using the NanoString GeoMx Digital Spatial Profiling WTA assay. A total of 32 glial regions of interest (ROIs) and 27 sarcomatous ROIs were analyzed. The ROIs chosen for analysis were selected using immunofluorescence for GFAP-rich (gliomatous) or GFAP-negative (sarcomatous) areas of viable tumor across at least 3 different areas of each tumor and represented a subset selected from 92 ROIs, after removing low-quality regions and regions that showed higher than expected GFAP or reticulin content upon analysis of serial stained sections (**Supplemental Figure S1**). Principal components analysis from transcriptomic data showed distinct grouping of sarcomatous ROIs, separate from the glial ROIs of the same individual’s tumor (**Figure 3A**). There were 3730 differentially expressed genes (DEGs) with an adjusted p-value <0.05 (**Figure 3B**). Gene set enrichment analysis (GSEA) of all DEGs identified upregulated pathways in sarcomatous regions such as extracellular matrix organization and degradation, ribosomal RNA processing, and the Mesenchymal Epithelial Transition (MET) signaling pathway (**Figure 3C**). The glial ROIs showed upregulated pathways involved in metal ion binding/response, neuronal projection development, and cellular signaling across synapses. Overall, GSEA supported a transcriptomic difference between glial and sarcomatous regions that aligns with a loss of central nervous system tissue processes/pathways and a gain of features related to mesenchymal tissue identity and generation/turnover of the extracellular matrix.

**Figure 3:**
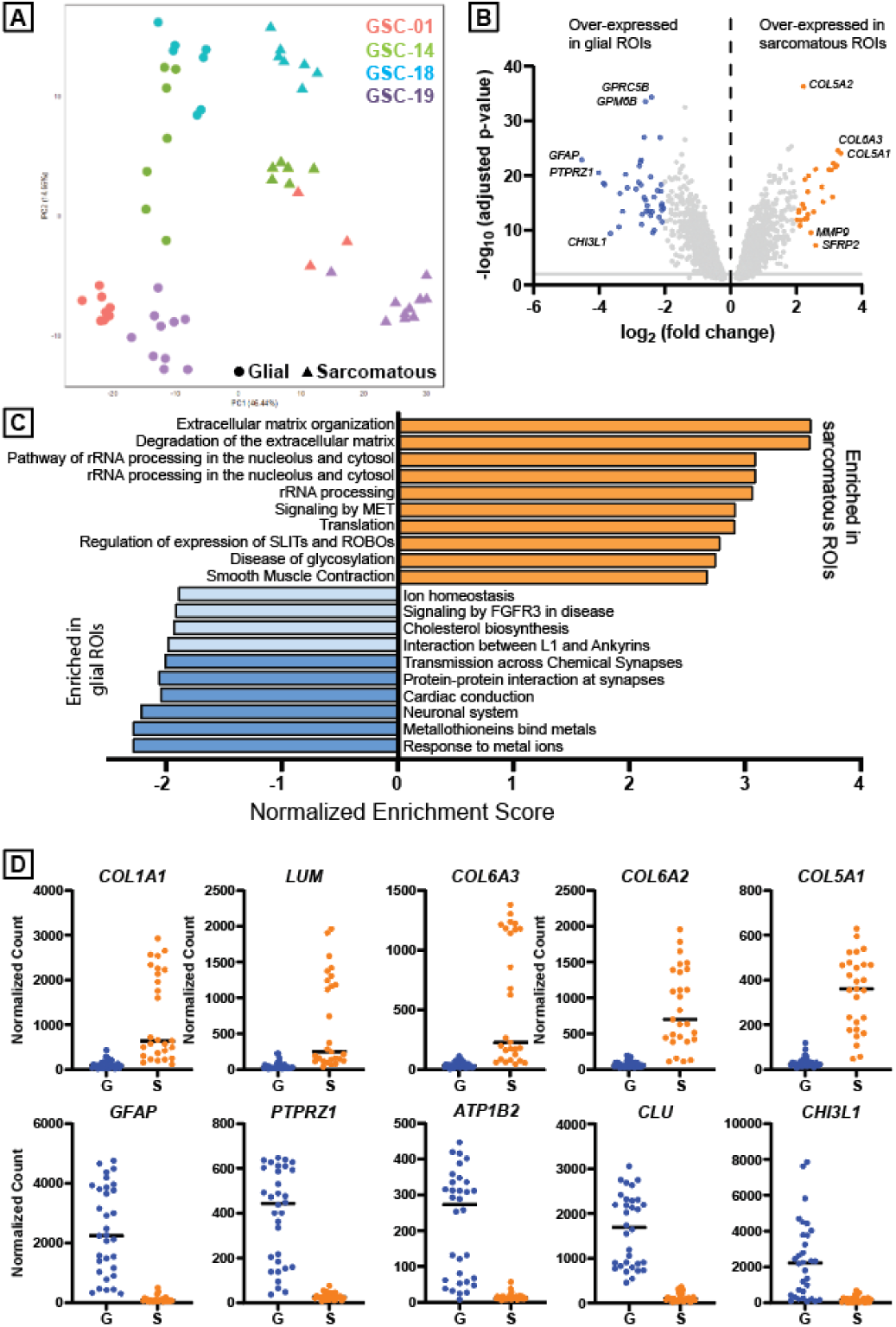
Digital spatial transcriptomic analysis of four gliosarcomas. (A) Principal components analysis shows separation of glial ROIs (circles) from sarcomatous ROIs (triangles) for all four individual tumors. (B) Volcano plot of differentially expressed genes between glial and sarcomatous ROIs. Colored dots represent 4-fold over-expression (orange) or under-expression (blue) in sarcomatous ROIs. (C) GSEA analysis of sarcomatous vs. glial ROIs. Darker shaded bars represent false discovery rate (FDR) ≤0.05. (D) Top five most over-expressed (top row) or under-expressed (bottom row) genes in sarcomatous (orange; “S”) versus glial (blue; “G”) ROIs.

To translate the DSP results back to histology, we applied a strict 4-fold change cutoff to the transcriptome data and identified 30 highly up-regulated DEGs and 42 highly down-regulated DEGs in sarcomatous ROIs. To account for variability between the 4 gliosarcomas (such as patient age, molecular features, tumor location, and age of material), we re-analyzed these 72 candidates in each individual tumor and observed that 24 of 30 (80%) of up-regulated DEGs and 37 of 42 (88%) of highly down-regulated DEGs were statistically significant in 2 or more individual gliosarcomas, lending confidence to the findings of the combined analysis (**Supplemental Figure S2**). The top 5 up-regulated genes in sarcomatous regions were *COL1A1*, *LUM*, *COL6A3*, *COL6A2*, and *COL5A1* (**Figure 3D**, top row). Other up-regulated genes in sarcomatous ROIs related to signaling pathways included: *SFRP2*, a secreted wingless (WNT) pathway modulatory protein (log_2_FC = 2.59, p <0.0001); *IGFBP4*, an insulin-like growth factor binding protein (log_2_FC = 2.19, p <0.0001); *CCN2*, encoding connective tissue growth factor (CTGF; log_2_FC = 2.53, p <0.0001); and *PDGFRB*, encoding platelet-derived growth factor receptor B (log_2_FC = 2.08, p <0.0001). The top 5 down-regulated genes in sarcomatous regions were *GFAP*, *PTPRZ1*, *ATP1B2*, *CLU*, and *CHI3L1* (**Figure 3D**, bottom row). *GFAP, PTPRZ1, ATPB2,* and *CLU* are associated with the astrocyte-like cellular state in glioblastoma (29). *CHI3L1* expression, in contrast, is associated with a mesenchymal-like cellular state in GBM (6, 29, 38). Reasoning that the mesenchymal regions of gliosarcomas may be an extreme form of the mesenchymal-like cellular state, we were intrigued to see downregulation of this marker in sarcomatous ROIs by transcription analysis. Therefore, we selected CHI3L1 for follow-up in tissue as a candidate down-regulated protein in sarcomatous regions.

### 3.4 CHI3L1 is differentially expressed in the glial and sarcomatous elements of most gliosarcomas

To evaluate protein expression level and pattern, immunohistochemistry for CHI3L1 was performed on 22 gliosarcomas and compared with staining pattern in an equal number of cases selected randomly from the GBM cohort. We noted labeling consistent with reactive astrocytes in infiltrated brain tissue, which is in accordance with published literature on CHI3L1 playing a role in the astrocytic response to injury (42). Consequently, for glioblastomas the most highly cellular regions of viable tumor were first identified by H&E and the same regions were cross-referenced to the CHI3L1 immunohistochemical stain for interpretation.

CHI3L1 immunoreactivity was low to moderate in most GBM cases, corresponding to 0 to 1+ intensity in about 75% of cases (**Figure 4A**). In contrast, the glial regions of GSC showed moderate to strong reactivity corresponding to 2+ to 3+ intensity in about 85% of cases and overlapped with GFAP-positive/reticulin-negative glial components (**Figure 4B**). Only 2 gliosarcomas were negative for CHI3L1, compared to 8 CHI3L1-negative glioblastomas (14% versus 32%, respectively – **Figure 4C**). In most gliosarcomas, CHI3L1 expression was reduced in the sarcomatous regions compared to glial regions within the same tumor, concordant with the transcriptomic findings (**Figure 4D**). Loss of CHI3L1 in sarcomatous regions of GSC could be related to a general loss of glial/astrocytic identity, akin to the reduction or loss of other glial markers like GFAP. Interestingly, two of 22 gliosarcomas showed strong CHI3L1 labeling in both glial and sarcomatous elements, suggesting that CHI3L1 protein expression can be uncoupled from GFAP expression in mesenchymal metaplasia in some cases.

**Figure 4:**
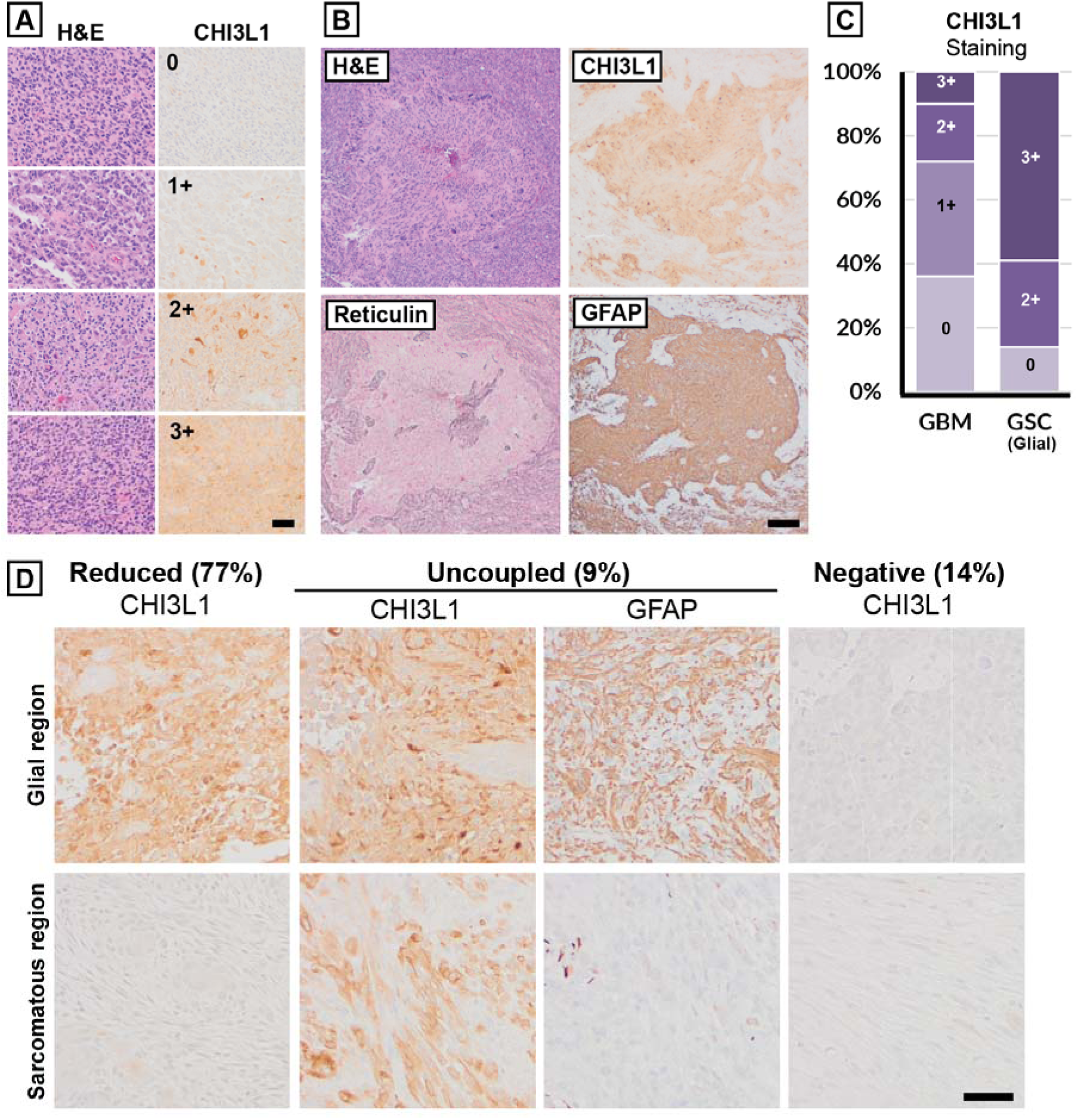
Immunohistochemical expression of CHI3L1. (A) Scoring system for CHI3L1 staining in glioblastoma. (B) Example of CHI3L1 reactivity in gliosarcoma, with labeling of glial (GFAP-high reticulin-low) but not sarcomatous (GFAP-low reticulin-high) elements. (C) Proportions of glioblastoma (left bar) or glial regions of gliosarcoma (right bar) staining at each intensity level. (D) Patterns of CHI3L1 staining in gliosarcomas; see text for details. Scale bars 50 μm (A, D) and 200 μm (B).

### 3.5 Sarcomatous elements of gliosarcoma show nuclear expression of the WNT effector protein LEF1

Identification of SFRP2 as a differentially expressed transcript was of interest due to the role of the WNT signaling pathway in the epithelial-mesenchymal transition (EMT) in GBM (20). Nuclear β-catenin labeling has been reported in sarcomatous regions in a subset of gliosarcomas and there is *in vitro* evidence that *SFRP2* over-expression can modulate a glial to mesenchymal transcriptomic shift in GBM tissue culture (13, 37). However, immunohistochemistry for β -catenin was negative in all four of the GSC cases used in transcriptome analysis (**Supplemental Figure S3**) and WTA showed only a marginal increase in *CTNNB1* transcript (Log_2_FC 0.3, p = 0.025). We attempted immunohistochemical analysis of SFRP2 but deemed it technically inadequate for interpretation (data not shown). Lymphoid-enhancer-binding factor 1 (LEF1) is a transcription factor that acts in a β-catenin-dependent or -independent manner depending on the context of cell type and intracellular signaling pathway activation, and can cooperate with an EMT transcription factor, zinc finger E-box binding homeobox 1 (Zeb1) in glioblastoma (17, 36). *LEF1* transcript was detected at a higher level in sarcomatous ROIs compared to glial ROIs in the DSP data (Log_2_FC = 0.8, adjusted p-value < 0.0001). Given evidence that LEF1 is a more sensitive histochemical marker for WNT-activated medulloblastoma than β-catenin, and that nuclear immunohistochemical expression of β -catenin is uncommon in GSC in a previously published study (13%; Schwetye *et al*), we pursued LEF1 for follow-up in the GSC cohort (1, 37). Nuclear LEF1 immunohistochemical labeling was observed focally in the endothelial cells within both GBM (**Figure 5A**) and in glial and sarcomatous regions of GSC. The glial elements of GSC cases were either negative for LEF1 nuclear reactivity (16 of 21 cases; 76%) or showed weak labeling of scattered glial/astrocytic cells (5/21 cases; 24%). In contrast, 20 of 21 GSC cases (95%) showed moderate to strong LEF1 nuclear labeling in at least 10% of the sarcomatous component, clearly distinct from labeling of the endothelial cells of microvascular proliferation (**Figure 5B**). Only 1 GSC was negative for LEF1 in the sarcomatous component. Overall, transcriptomic and immunohistochemical analysis indicates that nuclear LEF1 protein is at least focally to regionally increased in the sarcomatous elements of most gliosarcomas.

**Figure 5:**
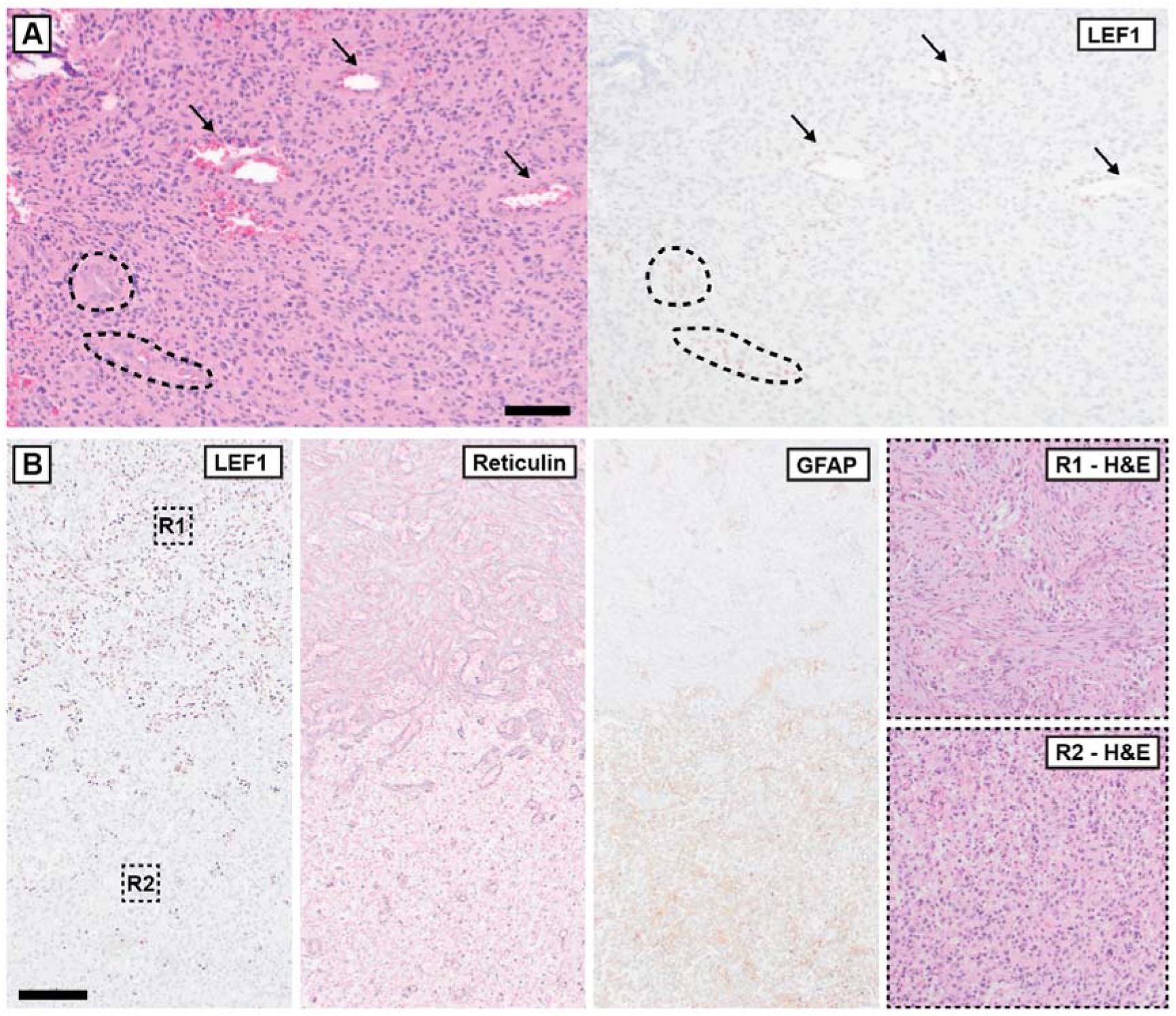
LEF1 immunohistochemistry. (A) Representative LEF1 staining in glioblastoma. Nuclear reactivity is restricted to vascular endothelial cells (black arrows) including in foci of early microvascular proliferation (dashed lines). (B) LEF1 staining in gliosarcoma. Nuclear reactivity is present in sarcomatous elements, overlapping with GFAP-negative reticulin-rich areas. The glial elements corresponding to GFAP-positive reticulin-poor regions have reactivity only in endothelial cells. Scale bars 100 μm (A) and 200 μm (B, applies to LEF1, reticulin, and GFAP).

## 4 Discussion

The GSC histologic subtype does not currently alter management for GBM patients, despite about a century of recognition. A better understanding of the features that set GSC apart from traditional GBM and identification of potential mechanisms and consequences of sarcomatous transformation in GSC are needed to help improve outcomes for patients. Studies on other GBM subtypes and histologic variants have led to meaningful discoveries about their underpinnings and diagnostic/prognostic/therapeutic implications. For instance, epithelioid GBM is known to have a high frequency of *BRAF* p.V600E mutation, giant cell GBM has an association with somatic or germline mismatch repair deficient state, and GBM with primitive neuronal component (GBM-PNC) is shown to have a higher rate of cerebrospinal fluid dissemination. Genomic, spatial transcriptomic, and proteomic analyses are powerful tools in the study of heterogeneous CNS tumors at a cellular and histologic level. For example, spatial transcriptomic analysis was recently used to implicate MYC signaling in the development of GBM-PNC (32). In the case of gliosarcomas, large cohorts examined in two recent studies showed evidence for a recurrent genetic pattern, following from earlier work showing low rates of EGFR activation in this subtype (9, 23, 34). Our study adds a third, independent NGS-based series demonstrating that primary adult gliosarcomas mostly lack *EGFR* alterations and are enriched for *NF1*, *TP53*, and PI3K pathway alterations, suggesting that a certain molecular background may be more permissive to sarcomatous transformation in this tumor at initial presentation.

GBM and, by extension, GSC are highly heterogeneous at the tissue and single-cell level. Bulk tumor gene expression analysis initially showed at least 3 transcriptomic subgroups, and more recent single-nucleus RNA sequencing studies have suggested at least 7, and possibly up to 10, different transcriptomically-defined GBM cellular states (31). The so-called mesenchymal-like (MES-like) state has long been noted to be enriched for *EGFR*-wildtype *NF1*-altered glioblastomas. Thus, the genetic features of our GSC series and other recent studies suggests a relationship between the MES-like glioblastoma cell state and the development of sarcomatous regions. However, it is unlikely that any particular glioma cell state or pattern of molecular alterations is either necessary or sufficient for GSC development, as there are ready examples of *EGFR*-activated gliosarcoma, as well as *EGFR*-unaltered/*NF1*-altered non-sarcomatous GBM. It is more plausible that a particular cellular state and/or genetic milieu is more permissive for GSC development. Further work is needed to identify the precise factors that control the sarcomatous phenotypic switch. It is interesting that the recently identified entity of oligosarcoma (IDH-mutant in most cases) shows a high rate of *NF1* alterations, as does desmoplastic melanoma - a tumor of neural crest origin with sarcoma-like histology (39, 40). A recent case study showed evidence for acquired *NF1* and *KRAS* mutations associated with sarcomatous transformation of a *BRAF* V600E-mutant GBM, suggesting a role for *NF1* alteration in sarcomatous transformation (30). Connections between *NF1* loss-of-function and mesenchymal phenotypic transition in glioblastoma have been suggested in at least one previous study (43).

We leveraged spatial transcriptomic results to evaluate CHI3L1, a marker of growing interest in neuro-oncology, and found evidence for a distinct expression pattern of this protein in gliosarcoma. CHI3L1 is an unusual protein that is implicated in neoplasia, inflammatory conditions, and neurodegenerative disease (25). CHI3L1 was recently identified as a marker and potential orchestrator of glioblastoma tumor cell connectivity which was associated with resistance to therapy and worse patient outcome (14). Therapies directed toward CHI3L1 are being explored and evaluation of these modalities in GSC may be warranted (8, 26). From a GSC phenotypic standpoint, least three main possibilities may be considered: first, CHI3L1 expression in GSC could indicate a particular cellular state within the glial regions, such as the mesenchymal-like state of which CHI3L1 is a marker; second, higher expression of CHI3L1 may be a response to the mesenchymal elements of the tumor, reflecting a change in astrocytic elements due to altered tumor microenvironment; third, CHI3L1 expression in malignant glial elements may have an active role in promoting sarcomatous transformation which subsequently down-regulate CHI3L1 expression during the phenotypic transition. Mechanistic studies would be necessary to differentiate these possibilities.

We note prior studies indicating transforming growth factor-β (TGF-β) signaling associated with histological transition from GBM to GSC and implicating nuclear factor-kappaB (NF-κB) in the mesenchymal-like state of GBM (4, 21). A few statistically significant changes in TGF-β family members were detected in our study and GSEA supported TGF-β pathway upregulation, while tumor necrosis factor alpha via NF-κB was balanced on GSEA (**Supplemental Figure S4**). The absence of a NF-κB pathway differential could be due to our DSP study design to detect differences between glial and mesenchymal elements at a single time point. If there is a high level of pathway activity in *both* the glial and mesenchymal elements of these tumors, there is in effect no “delta” to be uncovered by our approach. Our study should not be taken as narrowing the scope of pathways of interest in GSC, and our findings do not necessarily apply to GSC evolution over time and/or under treatment effects. This does raise the question of how sarcomatous transformation in GSC relates to, and may be different from, the more well-studied proneural-to-mesenchymal cellular state transition in GBM. Notably, a previous study on an epithelial-to-mesenchymal transition in glioma cancer stem cell models implicated a β-catenin-independent role for LEF1 in cooperation with zinc finger E-box-binding homeobox 1 (Zeb1) (36). High-confidence Zeb1 targets in that study were not differentially expressed in our analysis (data not shown).

Our study has limitations. It is a single-institution analysis that may be subject to bias in the pathological diagnosis of GSC. The GBM cohort was selected from cases that met the histological definition of the entity, and did not include cases that were diagnosed based on molecular features. We do not have detailed treatment histories to determine if different therapeutic approaches were used, which may have impacted outcome. Our spatial WTA approach is not at single-cell resolution; contributions from the non-neoplastic tumor microenvironment are not accounted for. The WTA cases were all from male patients and 3 of 4 cases were located in the temporal lobe, so generalizability of our findings may be limited. We did not perform epigenetic analysis of these tumors to further support their histological classification by a concordant DNA methylation family, subfamily, or class. While the molecular features in our series were, for the most part, concordant to IDH-wildtype glioblastoma, it is possible that some of our cases represent other, rare gliomas that overlap histologically and molecularly with GSC: for example, some histological gliosarcomas could fall into the methylation class of pleomorphic xanthoastrocytoma which was recently shown to encompass tumors that have GBM-like histology and shared molecular features (10). The cases selected for WTA varied in the age of the tissue. Although all ROIs in this analysis passed quality metrics, the number of DEGs varied in different cases and sensitivity for DEGs was lower in older cases. It was not feasible to study all of the potentially relevant pathways or DEGs; for example, our data could suggest consideration of the MET and CTGF-related pathways (among others) as contributors to GSC pathogenesis but were beyond our scope of study.

In summary, in this single-institution GSC cohort we confirm similar demographics and overall survival to GBM, suggest that a certain constellation of molecular features may predispose to this histologic subtype, demonstrate feasibility of spatial transcriptomic analysis in GSC, and suggest CHI3L1 and LEF1 (among other proteins) as drivers in GSC pathogenesis. Our results align with the findings of other recent series on the molecular features of GSC and additionally identify transcriptomic differences between sarcomatous and glial elements of these tumors which are correlated to differences in protein expression. Expression of CHI3L1 in glial elements of GSC may warrant further study to determine if this protein as a biomarker of, or an active regulator in, mesenchymal metaplasia in GBM. Further analysis of GSC using higher resolution spatial transcriptomic and proteomic analysis methods, potentially analyzing regional differences in DNA methylation and incorporating an analysis of the microenvironment, may be productive for further study.

## Author Contributions

MDW conceptualized the study, assembled the study cohorts, interpreted next-generation sequencing results, evaluated histology and immunohistochemistry, wrote the manuscript, and prepared figures. GZ and JL performed NanoString GeoMX Digital Spatial Profiling and analyzed the spatial transcriptomic data. TN performed the next-generation sequencing assay. KZ performed immunohistochemical studies. CC obtained funding for the study. All authors reviewed the manuscript.

## Acknowledgements

We thank Dr. Randall Woltjer and Ms. Victoria Krajbich, OHSU Department of Pathology and Laboratory Medicine, for performing CHI3L1 immunohistochemistry. We thank Dr. Monika Davare, OHSU School of Medicine, and Dr. Gregory Baker, OHSU Knight Cancer Institute, for discussion and feedback on the manuscript. Dr. Calixto-Hope Lucas provided data from reference (23).

## Funding Information

This study was supported by the Knight Cancer institute at Oregon Health & Science University and by the OHSU Department of Pathology and Laboratory Medicine.

## Conflict of Interest Statement

The authors declare that they have no conflicts of interest.

## Ethics Statement

This study was approved by the OHSU Internal Review Board with a waiver of patient consent (STUDY00027845 ).

---

**Table 1.** Frequencies of genetic alterations in GBM and GSC across multiple studies and data sets. Tumor type indicates glioblastoma (GBM) or gliosarcoma (GSC). For Lucas *et al.* data are from the initial tumor sampling for cases that maintained GBM histology at recurrence or transformed to GSC.

| Cohort | Tumor type | Number | *EGFR*-unaltered<br>*NF1*-altered | *EGFR*-unaltered<br>*NF1*-altered<br>*TP53*-altered | *EGFR*-Unaltered<br>*NF1*-Altered<br>*TP53*-Altered<br>*PI3K*-Altered |
| --- | --- | --- | --- | --- | --- |
| TCGA | GBM | 250 | 21 (8.4%) | 7 (2.8%) | 2 (0.8%) |
| Lucas *et al* | Primary GBM to recurrent GBM | 84 | 9 (10.7%) | 2 (2.4%) | 2 (2.4%) |
| This study | GBM | 75 | 7 (9.3%) | 1 (1.3%) | 1 (1.3%) |
| **Total** | **GBM** | **409** | **37 (9.0%)** | **10 (2.4%)** | **5 (1.2%)** |
| TCGA | GSC | 10 | 3 (30.0%) | 1 (10.0%) | 1 (10.0%) |
| Lucas *et al* | Primary GBM to recurrent GSC | 17 | 10 (58.8%) | 4 (23.5%) | 4 (23.0%) |
| Chen *et al* | GSC | 38 | 15 (39.5%) | 8 (21.0%) | 4 (10.5%) |
| This study | GSC | 25 | 12 (48.0%) | 5 (20.0%) | 5 (20.0%) |
| **Total** | **GSC** | **90** | **40 (44.4%)** | **18 (20.0%)** | **14 (15.5%)** |

## Notes

### Competing Interest Statement

The authors have declared no competing interest.

## References

1. Aboubakr O, Métais A, Doz F, Saffroy R, Masliah-Planchon J, Hasty L, Beccaria K, Ayrault O, Dufour C, Varlet P, Tauziède-Espariat A. LEF-1 immunohistochemistry, a better diagnostic biomarker than β-catenin for medulloblastoma, WNT-activated subtyping. J Neuropathol Exp Neurol. 2024 Jan 19;83(2):136–138. doi: 10.1093/jnen/nlad104. PMID: 38237134.

2. Alhalabi KT, Stichel D, Sievers P, Peterziel H, Sommerkamp AC, Sturm D, Wittmann A, Sill M, Jäger N, Beck P, Pajtler KW, Snuderl M, Jour G, Delorenzo M, Martin AM, Levy A, Dalvi N, Hansford JR, Gottardo NG, Uro-Coste E, Maurage CA, Godfraind C, Vandenbos F, Pietsch T, Kramm C, Filippidou M, Kattamis A, Jones C, Øra I, Mikkelsen TS, Zapotocky M, Sumerauer D, Scheie D, McCabe M, Wesseling P, Tops BBJ, Kranendonk MEG, Karajannis MA, Bouvier N, Papaemmanuil E, Dohmen H, Acker T, von Hoff K, Schmid S, Miele E, Filipski K, Kitanovski L, Krskova L, Gojo J, Haberler C, Alvaro F, Ecker J, Selt F, Milde T, Witt O, Oehme I, Kool M, von Deimling A, Korshunov A, Pfister SM, Sahm F, Jones DTW. PATZ1 fusions define a novel molecularly distinct neuroepithelial tumor entity with a broad histological spectrum. Acta Neuropathol. 2021 Nov;142(5):841–857. doi: 10.1007/s00401-021-02354-8. Epub 2021 Aug 21. PMID: 34417833; PMCID: PMC8500868.

3. Beaumont TL, Kupsky WJ, Barger GR, Sloan AE. Gliosarcoma with multiple extracranial metastases: case report and review of the literature. J Neurooncol. 2007 May;83(1):39–46. doi: 10.1007/s11060-006-9295-x. Epub 2006 Dec 14. PMID: 17171442.

4. Bhat KPL, Balasubramaniyan V, Vaillant B, Ezhilarasan R, Hummelink K, Hollingsworth F, Wani K, Heathcock L, James JD, Goodman LD, Conroy S, Long L, Lelic N, Wang S, Gumin J, Raj D, Kodama Y, Raghunathan A, Olar A, Joshi K, Pelloski CE, Heimberger A, Kim SH, Cahill DP, Rao G, Den Dunnen WFA, Boddeke HWGM, Phillips HS, Nakano I, Lang FF, Colman H, Sulman EP, Aldape K. Mesenchymal differentiation mediated by NF-κB promotes radiation resistance in glioblastoma. Cancer Cell. 2013 Sep 9;24(3):331–46. doi: 10.1016/j.ccr.2013.08.001. Epub 2013 Aug 29. PMID: 23993863; PMCID: PMC3817560.

5. Biernat W, Aguzzi A, Sure U, Grant JW, Kleihues P, Hegi ME. Identical mutations of the p53 tumor suppressor gene in the gliomatous and the sarcomatous components of gliosarcomas suggest a common origin from glial cells. J Neuropathol Exp Neurol. 1995 Sep;54(5):651–6. doi: 10.1097/00005072-199509000-00006. PMID: 7666053.

6. Boerman RH, Anderl K, Herath J, Borell T, Johnson N, Schaeffer-Klein J, Kirchhof A, Raap AK, Scheithauer BW, Jenkins RB. The glial and mesenchymal elements of gliosarcomas share similar genetic alterations. J Neuropathol Exp Neurol. 1996 Sep;55(9):973–81. doi: 10.1097/00005072-199609000-00004. PMID: 8800093.

7. Cancer Genome Atlas Research Network. Comprehensive genomic characterization defines human glioblastoma genes and core pathways. Nature. 2008 Oct 23;455(7216):1061–8. doi: 10.1038/nature07385. Epub 2008 Sep 4. Erratum in: Nature. 2013 Feb 28;494(7438):506. PMID: 18772890; PMCID: PMC2671642.

8. Chang MC, Chen CT, Chiang PF, Chiang YC. The Role of Chitinase-3-like Protein-1 (YKL40) in the Therapy of Cancer and Other Chronic-Inflammation-Related Diseases. Pharmaceuticals (Basel). 2024 Feb 27;17(3):307. doi: 10.3390/ph17030307. PMID: 38543093; PMCID: PMC10976000.

9. Chen L, Rizk E, Sherief M, Chang M, Lucas CH, Bettegowda C, Croog V, Mukherjee D, Rincon-Torroella J, Kamson DO, Huang P, Holdhoff M, Schreck K. Molecular characterization of gliosarcoma reveals prognostic biomarkers and clinical parallels with glioblastoma. J Neurooncol. 2025 Jan;171(2):403–411. doi: 10.1007/s11060-024-04859-0. Epub 2024 Oct 30. PMID: 39476147; PMCID: PMC11695672.

10. Dampier CH, Shah N, Galbraith K, Ebrahimi A, Neto OLA, Abdullaev Z, Alexandrescu S, Andreiuolo F, Armstrong T, Baker T, Cathcart S, Chung HJ, Cimino PJ, Conway KS, Cotter J, Costa FD, Dazelle K, Etminam N, Ferman SE, Fernandes I, Ferrone CK, Gilani A, Gilbert M, Gregory J, Ketchum C, Lee HS, Lee I, Lopes MBS, Mao Q, Marshall MS, McCord M, Neill SG, Nirschl JJ, Ozer BH, Paulus W, Penas-Prado M, Prinz M, Pytel P, Quezado M, Raffeld M, Rajan S, Ratliff M, Reifenberger G, Robinson L, Schittenhelm J, Schrimpf D, Singh O, Thomas C, Thomas D, Thomas-Ogunniyi J, Toland A, Turakulov R, Vaubel R, Wadhwani N, Wu J, Giannini C, Snuderl M, Brandner S, von Deimling A, Aldape K. Molecular, histologic, and clinical characterization of methylation class pleomorphic xanthoastrocytoma: An analysis of 469 tumors. Neurooncol Adv. 2025 Jul 19;7(1):vdaf089. doi: 10.1093/noajnl/vdaf089. PMID: 40735274; PMCID: PMC12305539.

11. Elizarraras JM, Liao Y, Shi Z, Zhu Q, Pico AR, Zhang B. WebGestalt 2024: faster gene set analysis and new support for metabolomics and multi-omics. Nucleic Acids Res. 2024 Jul 5;52(W1):W415-W421. doi: 10.1093/nar/gkae456. PMID: 38808672; PMCID: PMC11223849.

12. Feigin IH, Gross SW. Sarcoma arising in glioblastoma of the brain. Am J Pathol. 1955 Jul-Aug;31(4):633-53. PMID: 14388124; PMCID: PMC1942557.

13. Guo M, Goudarzi KM, Abedi S, Pieber M, Sjöberg E, Behnan J, Zhang XM, Harris RA, Bartek J, Lindström MS, Nistér M, Hägerstrand D. SFRP2 induces a mesenchymal subtype transition by suppression of SOX2 in glioblastoma. Oncogene. 2021 Aug;40(32):5066–5080. doi: 10.1038/s41388-021-01825-2. Epub 2021 May 21. Erratum in: Oncogene. 2021 Aug;40(32):5153. doi: 10.1038/s41388-021-01929-9. PMID: 34021259; PMCID: PMC8363098.

14. Hai L, Hoffmann DC, Wagener RJ, Azorin DD, Hausmann D, Xie R, Huppertz MC, Hiblot J, Sievers P, Heuer S, Ito J, Cebulla G, Kourtesakis A, Kaulen LD, Ratliff M, Mandelbaum H, Jung E, Jabali A, Horschitz S, Ernst KJ, Reibold D, Warnken U, Venkataramani V, Will R, Suvà ML, Herold-Mende C, Sahm F, Winkler F, Schlesner M, Wick W, Kessler T. A clinically applicable connectivity signature for glioblastoma includes the tumor network driver CHI3L1. Nat Commun. 2024 Feb 6;15(1):968. doi: 10.1038/s41467-024-45067-8. PMID: 38320988; PMCID: PMC10847113.

15. Jacobsen J, Locallo A, O’Rourke CJ, Carlsen JF, Cohen J, Ewald JD, Scheie D, Grunnet K, Schmidt AY, Melchior LC, Larsen VA, Andersen JB, Poulsen HS, Weischenfeldt J, Broholm H, Michaelsen SR, Kristensen BW. Clinical, Radiological, and Molecular Insights into Extracranial Metastases from Adult Gliomas. Neuro Oncol. 2025 Aug 16:noaf178. doi: 10.1093/neuonc/noaf178. Epub ahead of print. PMID: 40842361.

16. Kim Y, Danaher P, Cimino PJ, Hurth K, Warren S, Glod J, Beechem JM, Zada G, McEachron TA. Highly Multiplexed Spatially Resolved Proteomic and Transcriptional Profiling of the Glioblastoma Microenvironment Using Archived Formalin-Fixed Paraffin-Embedded Specimens. Mod Pathol. 2023 Jan;36(1):100034. doi: 10.1016/j.modpat.2022.100034. PMID: 36788070; PMCID: PMC9937641.

17. Kobayashi W, Ozawa M. The epithelial-mesenchymal transition induced by transcription factor LEF-1 is independent of β-catenin. Biochem Biophys Rep. 2018 Jun 12;15:13–18. doi: 10.1016/j.bbrep.2018.06.003. PMID: 29998192; PMCID: PMC6038150.

18. La Torre D, Della Torre A, Lo Turco E, Longo P, Pugliese D, Lacroce P, Raudino G, Romano A, Lavano A, Tomasello F. Primary Intracranial Gliosarcoma: Is It Really a Variant of Glioblastoma? An Update of the Clinical, Radiological, and Biomolecular Characteristics. J Clin Med. 2023 Dec 22;13(1):83. doi: 10.3390/jcm13010083. PMID: 38202090; PMCID: PMC10779593.

19. Landvater R, Tripathy A, Nieblas-Bedolla E, Shao L, Conway K, Al-Holou W, Ferris SP. Sarcomatous transformation of IDH-mutant astrocytoma matching to methylation class oligosarcoma following embolization, a case report. Acta Neuropathol Commun. 2024 Dec 20;12(1):196. doi: 10.1186/s40478-024-01908-7. PMID: 39707480; PMCID: PMC11662476.

20. Lee Y, Lee JK, Ahn SH, Lee J, Nam DH. WNT signaling in glioblastoma and therapeutic opportunities. Lab Invest. 2016 Feb;96(2):137–50. doi: 10.1038/labinvest.2015.140. Epub 2015 Dec 7. PMID: 26641068.

21. Li A, Hancock JC, Quezado M, Ahn S, Briceno N, Celiku O, Ranjan S, Aboud O, Colwell N, Kim SA, Nduom E, Kuhn S, Park DM, Vera E, Aldape K, Armstrong TS, Gilbert MR. TGF-β and BMP signaling are associated with the transformation of glioblastoma to gliosarcoma and then osteosarcoma. Neurooncol Adv. 2023 Dec 19;6(1):vdad164. doi: 10.1093/noajnl/vdad164. PMID: 38292240; PMCID: PMC10825841.

22. Li MM, Datto M, Duncavage EJ, Kulkarni S, Lindeman NI, Roy S, Tsimberidou AM, Vnencak-Jones CL, Wolff DJ, Younes A, Nikiforova MN. Standards and Guidelines for the Interpretation and Reporting of Sequence Variants in Cancer: A Joint Consensus Recommendation of the Association for Molecular Pathology, American Society of Clinical Oncology, and College of American Pathologists. J Mol Diagn. 2017 Jan;19(1):4–23. doi: 10.1016/j.jmoldx.2016.10.002. PMID: 27993330; PMCID: PMC5707196.

23. Lucas CG, Al-Adli NN, Young JS, Gupta R, Morshed RA, Wu J, Ravindranathan A, Shai A, Oberheim Bush NA, Taylor JW, de Groot J, Villanueva-Meyer JE, Pekmezci M, Perry A, Bollen AW, Theodosopoulos PV, Aghi MK, Chang EF, Hervey-Jumper SL, Raleigh DR, Molinaro AM, Costello JF, Diaz AA, Clarke JL, Butowski NA, Phillips JJ, Chang SM, Berger MS, Solomon DA. Longitudinal multimodal profiling of IDH-wildtype glioblastoma reveals the molecular evolution and cellular phenotypes underlying prognostically different treatment responses. Neuro Oncol. 2025 Jan 12;27(1):89–105. doi: 10.1093/neuonc/noae214. PMID: 39560080; PMCID: PMC11726253.

24. Moffet JJD, Fatunla OE, Freytag L, Kriel J, Jones JJ, Roberts-Thomson SJ, Pavenko A, Scoville DK, Zhang L, Liang Y, Morokoff AP, Whittle JR, Freytag S, Best SA. Spatial architecture of high-grade glioma reveals tumor heterogeneity within distinct domains. Neurooncol Adv. 2023 Nov 1;5(1):vdad142. doi: 10.1093/noajnl/vdad142. PMID: 38077210; PMCID: PMC10699851.

25. Mwale PF, Hsieh CT, Yen TL, Jan JS, Taliyan R, Yang CH, Yang WB. Chitinase-3-like-1: a multifaceted player in neuroinflammation and degenerative pathologies with therapeutic implications. Mol Neurodegener. 2025 Jan 18;20(1):7. doi: 10.1186/s13024-025-00801-8. PMID: 39827337; PMCID: PMC11742494.

26. Nada H, Zhang L, Kaur B, Gabr MT. CHI3L1-targeted small molecules as glioblastoma therapies: Virtual screening-based discovery, biophysical validation, pharmacokinetic profiling, and evaluation in glioblastoma spheroids. Eur J Med Chem. 2025 Nov 5;297:117960. doi: 10.1016/j.ejmech.2025.117960. Epub 2025 Jul 8. PMID: 40663973.

27. Nagaishi M, Paulus W, Brokinkel B, Vital A, Tanaka Y, Nakazato Y, Giangaspero F, Ohgaki H. Transcriptional factors for epithelial-mesenchymal transition are associated with mesenchymal differentiation in gliosarcoma. Brain Pathol. 2012 Sep;22(5):670–6. doi: 10.1111/j.1750-3639.2012.00571.x. Epub 2012 Mar 5. PMID: 22288519; PMCID: PMC8057633.

28. Nagaishi M, Kim YH, Mittelbronn M, Giangaspero F, Paulus W, Brokinkel B, Vital A, Tanaka Y, Nakazato Y, Legras-Lachuer C, Lachuer J, Ohgaki H. Amplification of the STOML3, FREM2, and LHFP genes is associated with mesenchymal differentiation in gliosarcoma. Am J Pathol. 2012 May;180(5):1816–23. doi: 10.1016/j.ajpath.2012.01.027. PMID: 22538188.

29. Neftel C, Laffy J, Filbin MG, Hara T, Shore ME, Rahme GJ, Richman AR, Silverbush D, Shaw ML, Hebert CM, Dewitt J, Gritsch S, Perez EM, Gonzalez Castro LN, Lan X, Druck N, Rodman C, Dionne D, Kaplan A, Bertalan MS, Small J, Pelton K, Becker S, Bonal D, Nguyen QD, Servis RL, Fung JM, Mylvaganam R, Mayr L, Gojo J, Haberler C, Geyeregger R, Czech T, Slavc I, Nahed BV, Curry WT, Carter BS, Wakimoto H, Brastianos PK, Batchelor TT, Stemmer-Rachamimov A, Martinez-Lage M, Frosch MP, Stamenkovic I, Riggi N, Rheinbay E, Monje M, Rozenblatt-Rosen O, Cahill DP, Patel AP, Hunter T, Verma IM, Ligon KL, Louis DN, Regev A, Bernstein BE, Tirosh I, Suvà ML. An Integrative Model of Cellular States, Plasticity, and Genetics for Glioblastoma. Cell. 2019 Aug 8;178(4):835–849.e21. doi: 10.1016/j.cell.2019.06.024. Epub 2019 Jul 18. PMID: 31327527; PMCID: PMC6703186.

30. Nelson BE, Reddy NK, Huse JT, Amini B, Nardo M, Gouda M, Weathers SP, Subbiah V. Histological transformation to gliosarcoma with combined BRAF/MEK inhibition in BRAF V600E mutated glioblastoma. NPJ Precis Oncol. 2023 May 25;7(1):47. doi: 10.1038/s41698-023-00398-5. PMID: 37231247; PMCID: PMC10212928.

31. Nomura M, Spitzer A, Johnson KC, Garofano L, Nehar-Belaid D, Galili Darnell N, Greenwald AC, Bussema L, Oh YT, Varn FS, D’Angelo F, Gritsch S, Anderson KJ, Migliozzi S, Gonzalez Castro LN, ChowdhFury T, Robine N, Reeves C, Park JB, Lipsa A, Hertel F, Golebiewska A, Niclou SP, Nusrat L, Kellet S, Das S, Moon HE, Paek SH, Bielle F, Laurenge A, Di Stefano AL, Mathon B, Picca A, Sanson M, Tanaka S, Saito N, Ashley DM, Keir ST, Ligon KL, Huse JT, Yung WKA, Lasorella A, Verhaak RGW, Iavarone A, Suvà ML, Tirosh I. The multilayered transcriptional architecture of glioblastoma ecosystems. Nat Genet. 2025 May;57(5):1155–1167. doi: 10.1038/s41588-025-02167-5. Epub 2025 May 9. PMID: 40346361; PMCID: PMC12081307.

32. Pagani F, Orzan F, Lago S, De Bacco F, Prelli M, Cominelli M, Somenza E, Gryzik M, Balzarini P, Ceresa D, Marubbi D, Isella C, Crisafulli G, Poli M, Malatesta P, Galli R, Ronca R, Zippo A, Boccaccio C, Poliani PL. Concurrent RB1 and P53 pathway disruption predisposes to the development of a primitive neuronal component in high-grade gliomas depending on MYC-driven EBF3 transcription. Acta Neuropathol. 2025 Jan 16;149(1):8. doi: 10.1007/s00401-025-02845-y. PMID: 39821672.

33. Patel AP, Tirosh I, Trombetta JJ, Shalek AK, Gillespie SM, Wakimoto H, Cahill DP, Nahed BV, Curry WT, Martuza RL, Louis DN, Rozenblatt-Rosen O, Suvà ML, Regev A, Bernstein BE. Single-cell RNA-seq highlights intratumoral heterogeneity in primary glioblastoma. Science. 2014 Jun 20;344(6190):1396–401. doi: 10.1126/science.1254257. Epub 2014 Jun 12. PMID: 24925914; PMCID: PMC4123637.

34. Reis RM, Könü-Lebleblicioglu D, Lopes JM, Kleihues P, Ohgaki H. Genetic profile of gliosarcomas. Am J Pathol. 2000 Feb;156(2):425–32. doi: 10.1016/S0002-9440(10)64746-3. PMID: 10666371; PMCID: PMC1850048.

35. Rodriguez FJ, Scheithauer BW, Perry A, Oliveira AM, Jenkins RB, Oviedo A, Mork SJ, Palmer CA, Burger PC. Ependymal tumors with sarcomatous change (“ependymosarcoma”): a clinicopathologic and molecular cytogenetic study. Am J Surg Pathol. 2008 May;32(5):699–709. doi: 10.1097/PAS.0b013e318158234e. PMID: 18347506.

36. Rosmaninho P, Mükusch S, Piscopo V, Teixeira V, Raposo AA, Warta R, Bennewitz R, Tang Y, Herold-Mende C, Stifani S, Momma S, Castro DS. Zeb1 potentiates genome-wide gene transcription with Lef1 to promote glioblastoma cell invasion. EMBO J. 2018 Aug 1;37(15):e97115. doi: 10.15252/embj.201797115. Epub 2018 Jun 14. PMID: 29903919; PMCID: PMC6068449.

37. Schwetye KE, Joseph NM, Al-Kateb H, Rich KM, Schmidt RE, Perry A, Gutmann DH, Dahiya S. Gliosarcomas lack BRAF^V600E^ mutation, but a subset exhibit β-catenin nuclear localization. Neuropathology. 2016 Oct;36(5):448–455. doi: 10.1111/neup.12293. Epub 2016 Mar 2. PMID: 26932501.

38. Spitzer A, Johnson KC, Nomura M, Garofano L, Nehar-Belaid D, Darnell NG, Greenwald AC, Bussema L, Oh YT, Varn FS, D’Angelo F, Gritsch S, Anderson KJ, Migliozzi S, Gonzalez Castro LN, Chowdhury T, Robine N, Reeves C, Park JB, Lipsa A, Hertel F, Golebiewska A, Niclou SP, Nusrat L, Kellet S, Das S, Moon HE, Paek SH, Bielle F, Laurenge A, Di Stefano AL, Mathon B, Picca A, Sanson M, Tanaka S, Saito N, Ashley DM, Keir ST, Ligon KL, Huse JT, Yung WKA, Lasorella A, Iavarone A, Verhaak RGW, Tirosh I, Suvà ML. Deciphering the longitudinal trajectories of glioblastoma ecosystems by integrative single-cell genomics. Nat Genet. 2025 May;57(5):1168–1178. doi: 10.1038/s41588-025-02168-4. Epub 2025 May 9. PMID: 40346362; PMCID: PMC12081298.

39. Suwala AK, Felix M, Friedel D, Stichel D, Schrimpf D, Hinz F, Hewer E, Schweizer L, Dohmen H, Pohl U, Staszewski O, Korshunov A, Stein M, Wongsurawat T, Cheunsuacchon P, Sathornsumetee S, Koelsche C, Turner C, Le Rhun E, Mühlebner A, Schucht P, Özduman K, Ono T, Shimizu H, Prinz M, Acker T, Herold-Mende C, Kessler T, Wick W, Capper D, Wesseling P, Sahm F, von Deimling A, Hartmann C, Reuss DE. Oligosarcomas, IDH-mutant are distinct and aggressive. Acta Neuropathol. 2022 Feb;143(2):263–281. doi: 10.1007/s00401-021-02395-z. Epub 2021 Dec 30. PMID: 34967922; PMCID: PMC8742817.

40. Wiesner T, Kiuru M, Scott SN, Arcila M, Halpern AC, Hollmann T, Berger MF, Busam KJ. NF1 Mutations Are Common in Desmoplastic Melanoma. Am J Surg Pathol. 2015 Oct;39(10):1357–62. doi: 10.1097/PAS.0000000000000451. PMID: 26076063; PMCID: PMC5037960.

41. WHO Classification of Tumours Editorial Board. World Health Organization classification of tumours of the central nervous system. 5th ed. Lyon: International Agency for Research on Cancer; 2021.

42. Wiley CA, Bonneh-Barkay D, Dixon CE, Lesniak A, Wang G, Bissel SJ, Kochanek PM. Role for mammalian chitinase 3-like protein 1 in traumatic brain injury. Neuropathology. 2015 Apr;35(2):95–106. doi: 10.1111/neup.12158. Epub 2014 Nov 7. PMID: 25377763.

43. Wood MD, Reis GF, Reuss DE, Phillips JJ. Protein Analysis of Glioblastoma Primary and Posttreatment Pairs Suggests a Mesenchymal Shift at Recurrence. J Neuropathol Exp Neurol. 2016 Oct;75(10):925–935. doi: 10.1093/jnen/nlw068. Epub 2016 Aug 18. PMID: 27539476; PMCID: PMC6281078.

44. Wood MD, Beadling C, Neff T, Moore S, Harrington CA, Baird L, Corless C. Molecular profiling of pre- and post-treatment pediatric high-grade astrocytomas reveals acquired increased tumor mutation burden in a subset of recurrences. Acta Neuropathol Commun. 2023 Sep 5;11(1):143. doi: 10.1186/s40478-023-01644-4. PMID: 37670377; PMCID: PMC10481558.

45. Zaki MM, Mashouf LA, Woodward E, Langat P, Gupta S, Dunn IF, Wen PY, Nahed BV, Bi WL. Genomic landscape of gliosarcoma: distinguishing features and targetable alterations. Sci Rep. 2021 Sep 9;11(1):18009. doi: 10.1038/s41598-021-97454-6. PMID: 34504233; PMCID: PMC8429571.

46. Zhang B, Kirov S, Snoddy J. WebGestalt: an integrated system for exploring gene sets in various biological contexts. Nucleic Acids Res. 2005 Jul 1;33(Web Server issue):W741-8. doi: 10.1093/nar/gki475. PMID: 15980575; PMCID: PMC1160236.

