## Supplemental Text and Figures for "Clinical and molecular features of primary gliosarcoma with digital spatial whole-transcriptome analysis of glial and mesenchymal components"

Supplemental Figure S1

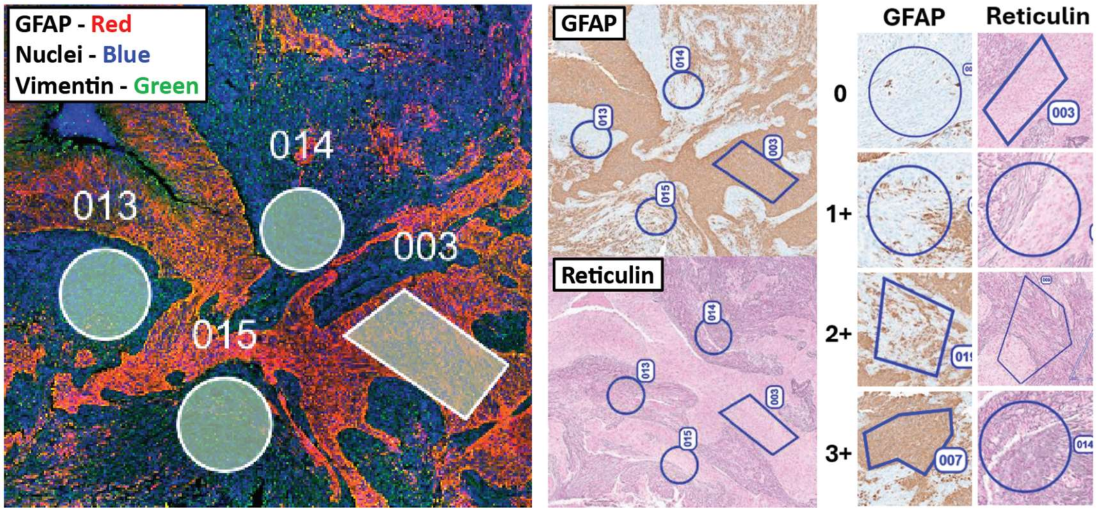


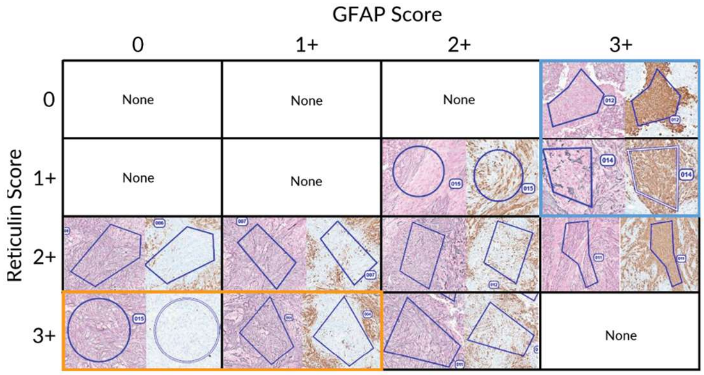


| | | | |
|---|---|---|---|
| **0** | **0** | **0** | **18** |
| 0 | 0 | 2 | 17 |
| 1 | 2 | 9 | 11 |
| 9 | 19 | 7 | 0 |

| Case | Age | Sex | Location | Glial ROI | Mesen. ROI | Genetic alterations |
|---|---|---|---|---|---|---|
| GSC-01 | 65 | M | Temporal | 8 | 3 | *TERT*p, *TP53*, *NF1*, *PTEN*, +7/-10, MGMT Methylated |
| GSC-14 | 61 | M | Temporal | 7 | 6 | *TERT*p, *TP53*, *PTEN*, *PIK3R1*, *RB1*, MGMT Not methylated |
| GSC-18 | 78 | M | Frontal | 7 | 7 | *TERT*p, *NF1*, *PTEN*, MGMT Unknown |
| GSC-19 | 69 | M | Temporal | 19 | 11 | *TERT*p, *TP53*, *PTEN*, MGMT Unknown |
| **TOTALS** | | | | **32** | **27** | |

Supplemental Figure S1 – Selection of ROIs for spatial transcriptomic analysis. Regions of approximately 100 to 500 nucleated cells were selected under immunofluorescence for GFAP and vimentin with Syto-13 counter-staining. ROIs were manually translated to consecutive H&E, GFAP, and reticulin stained sections and each region was scored for the degree of GFAP reactivity and reticulin deposition. The H&E sections were used for morphologic correlation. A matrix of highest differentially-labeled regions was generated and highest quality regions were determined as described in the main text. Three glial ROIs and one sarcomatous ROI in the blue or orange cells respectively were excluded due to poor correlation with morphology by H&E. The final tally of ROIs and corresponding genetic and clinical features are tabulated.

Supplemental Figure S2

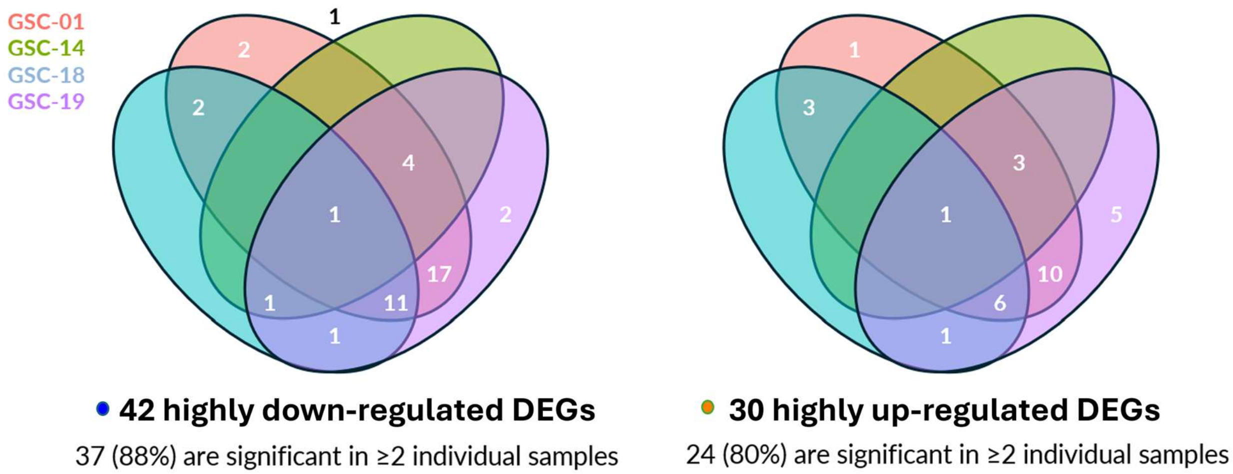


Supplemental Figure S2 – DEGs were identified by statistical significance level and by 4-fold up-regulation in sarcomatous ROIs (right) or 4-fold down-regulation in sarcomatous ROIs. Since the DEGs were identified in a combined analysis of 4 different tumors, these data represent how many DEGs were significant in the four tumors when analyzed individually.

Supplemental Figure S3

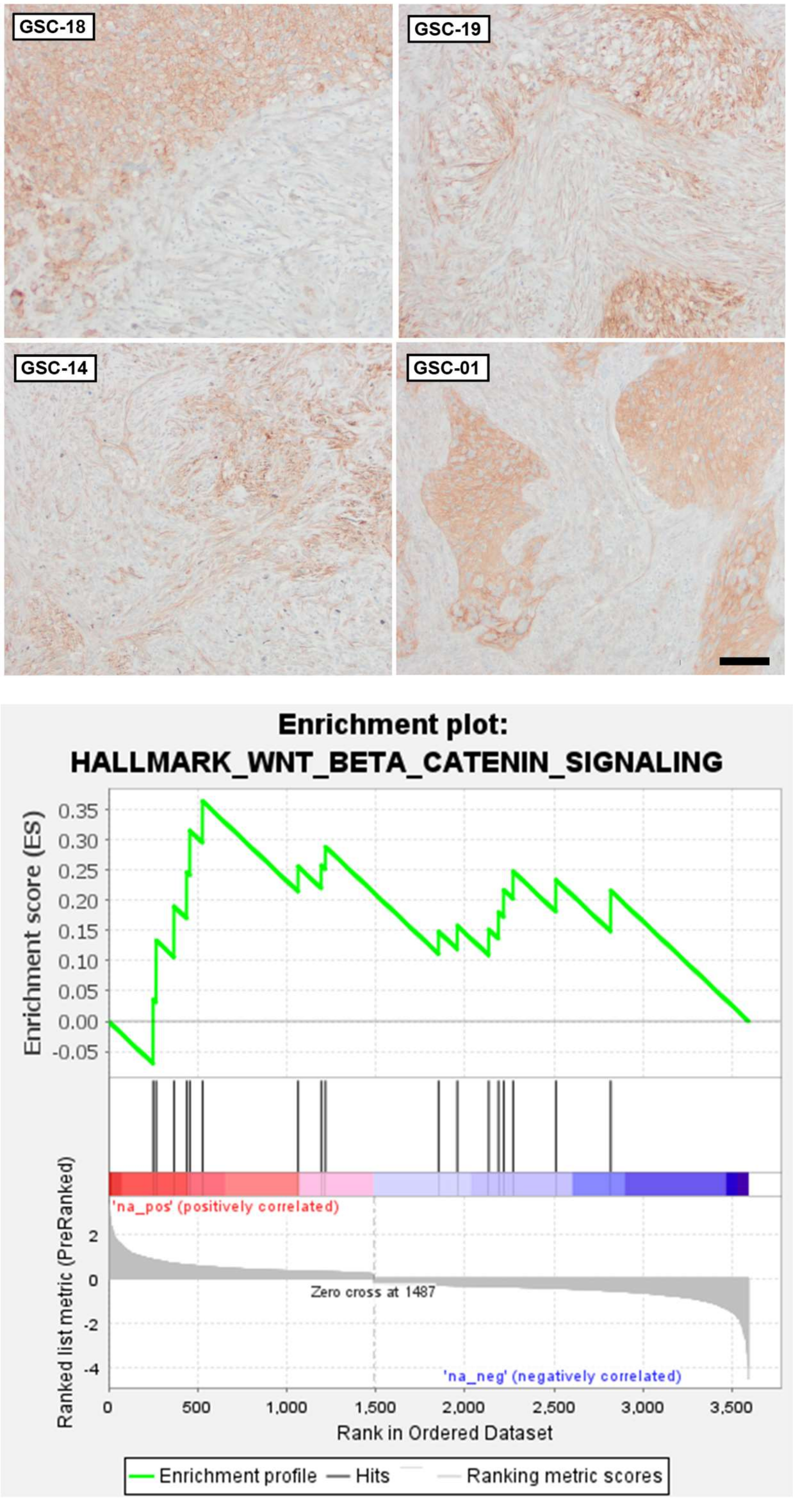


Supplemental Figure S3: B-catenin immunohistochemistry in four gliosarcoma cases used for DSP analysis. Scale bar 100 μm. Enrichment plot for WNT-b-catenin on GSEA using the HALLMARK gene sets for analysis.

Supplemental Figure S4

<u>Hallmark TGFb Signaling</u>

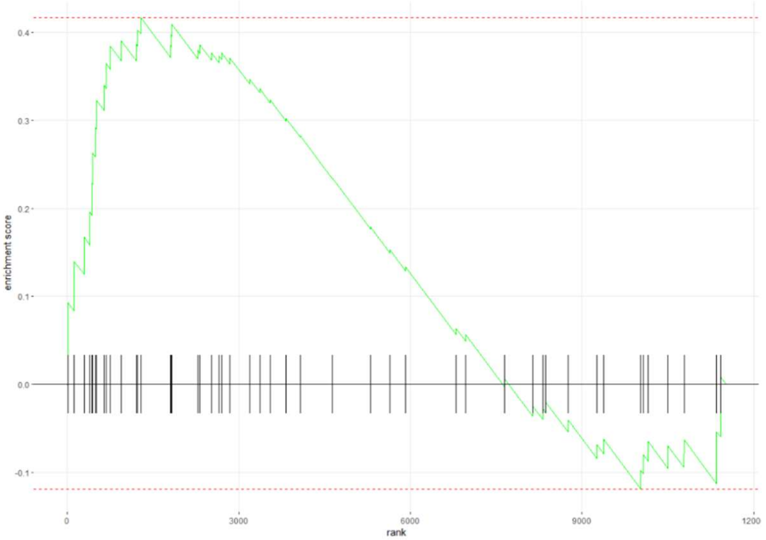


| Gene | Log2FC | p-value |
|---|---|---|
| TGFB1 | 0.5 | <0.0001 |
| TGFB1\|1 | 0.8 | <0.0001 |
| TGFB2 | -0.9 | <0.0001 |
| TGFB3 | 0.7 | <0.0001 |
| TGFBI | 2.3 | <0.0001 |
| TGFBR1 | 0.5 | <0.0001 |

<u>Hallmark TNFA Signaling by NFKB</u>

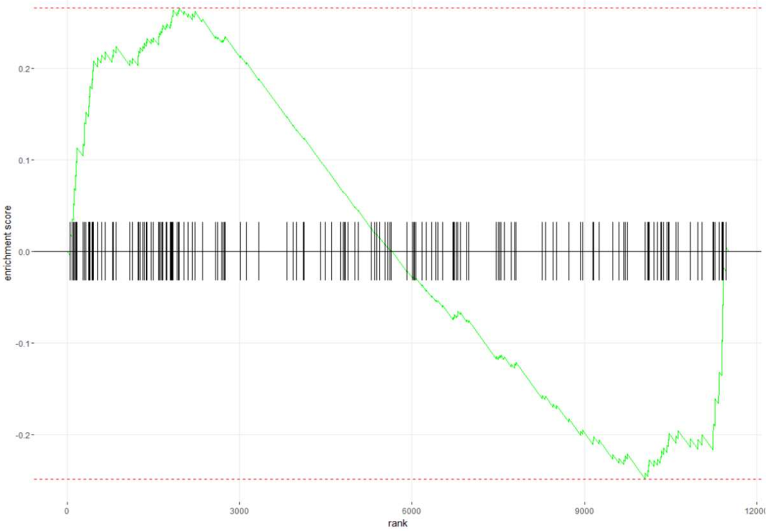


| Gene | Log2FC | p-value |
|---|---|---|
| NFKB1 | 0.3 | 0.031 |
| NFKB2 | -0.3. | 0.041 |

Supplemental Figure S4. HALLMARK enrichment plots for TGF-b signaling (upper) and for TNFA-related TGF-b signaling (lower), with differential expression of related genes from GSEA given in adjacent tables.
