## Supplemental Table 1 for "Clinical and molecular features of primary gliosarcoma with digital spatial whole-transcriptome analysis of glial and mesenchymal components"

| Case | Gene | Transcript | cDNA Variant | Protein Variant | Coordinates | Reference | Variant | VAF | Interp. |
|---|---|---|---|---|---|---|---|---|---|
| GSC-01 | TP53 | NM_000546 | c.818G>A | p.R273H | chr17:7577120 | C | T | 0.43 | P |
| | TERT | CCDS3861.2 | overlaps 1-kb region upstream | Promoter region | chr5:1295228 | G | A | 0.34 | P |
| | PTEN | CCDS31238.1 | c.402G>C | p.M134I | chr10:89692918 | G | C | 0.43 | P |
| | NF1 | CCDS42292.1 | c.5907_5908del | p.R1970fs*6 | chr17:29661948 | CAA | C | 0.48 | P |
| GSC-02 | TERT | CCDS3861.2 | overlaps 1-kb region upstream | Promoter region | chr5:1295228 | G | A | 0.38 | P |
| | TP53 | NM_000546 | c.764_766del | p.I255del | chr17:7577514 | GTGA | G | 0.55 | P |
| | NF1 | CCDS42292.1 | c.3144G>A | p.W1048* | chr17:29557890 | G | A | 0.26 | P |
| | PIK3R1 | CCDS3993.1 | c.1686_1691del | p.M563_N564del | chr5:67591092 | GTATGAA | G | 0.39 | P |
| GSC-03 | KDR | CCDS3497.1 | c.3064C>T | p.R1022* | chr4:55958789 | G | A | 0.2 | LP |
| | NBN | CCDS6249.1 | c.2140C>T | p.R714* | chr8:90955525 | G | A | 0.27 | P |
| | PIK3R1 | CCDS3993.1 | c.1002_1005del | p.Y334fs*1 | chr5:67588171 | ACTGG | A | 0.28 | P |
| | PTEN | CCDS31238.1 | c.419T>G | p.L140* | chr10:89692935 | T | G | 0.3 | P |
| | NF1 | CCDS42292.1 | c.7226C>G | p.T2409R | chr17:29676174 | C | G | 0.39 | LP |
| GSC-04 | TP53 | NM_000546 | c.527G>A | p.C176Y | chr17:7578403 | C | T | 0.31 | P |
| | TERT | CCDS3861.2 | overlaps 1-kb region upstream | Promoter region | chr5:1295228 | G | A | 0.41 | P |
| | NOTCH 1 | CCDS43905.1 | c.3826T>C | p.C1276R | chr9:139401243 | A | G | 0.5 | VUS |
| GSC-05 | ALK | CCDS33172.1 | c.4200A>C | p.E1400D | chr2:29416753 | T | G | 0.52 | VUS |
| | TERT | CCDS3861.2 | overlaps 1-kb region upstream | Promoter region | chr5:1295228 | G | A | 0.56 | P |
| | PTEN | CCDS31238.1 | c.521A>G | p.Y174C | chr10:89711903 | A | G | 0.78 | LP |
| | NF1 | CCDS42292.1 | c.305_306del | p.M102fs*4 | chr17:29490219 | ATG | A | 0.44 | LP |
| | NF1 | CCDS42292.1 | c.3622delT | p.L1208fs*7 | chr17:29560144 | AT | A | 0.42 | LP |
| GSC-06 | TP53 | NM_000546 | c.818G>A | p.R273H | chr17:7577120 | C | T | 0.54 | P |
| | PTEN | CCDS31238.1 | c.98T>G | p.I33S | chr10:89653800 | T | G | 0.48 | P |
| | TERT | CCDS3861.2 | overlaps 1-kb region upstream | Promoter region | chr5:1295228 | G | A | 0.28 | P |
| | RB1 | CCDS31973.1 | c.1064_1065del | p.R355Nfs*6 | chr13:48942673 | CAG | C | 0.36 | P |
| | NF1 | CCDS42292.1 | c.4475delT | p.V1492fs*8 | chr17:29587430 | GT | G | 0.48 | LP |
| GSC-07 | TP53 | NM_000546 | c.376-1G>A | Splice site | chr17:7578555 | C | T | 0.6 | P |
| | PIK3R1 | CCDS3993.1 | c.1126G>A | p.G376R | chr5:67589138 | G | A | 0.27 | P |
| | PTEN | CCDS31238.1 | c.517C>T | p.R173C | chr10:89711899 | C | T | 0.35 | P |
| | TERT | CCDS3861.2 | overlaps 1-kb region upstream | Promoter region | chr5:1295228 | G | A | 0.46 | P |

| | | | | | | | | | |
|---|---|---|---|---|---|---|---|---|---|
| | SUFU | CCDS7537.1 | c.546C>A | p.D182E | chr10:104352430 | C | A | 0.5 | VUS |
| GSC-08 | TERT | CCDS3861.2 | overlaps 1-kb region upstream | Promoter region | chr5:1295228 | G | A | 0.54 | P |
| | EGFR | Multiple | | | chr7: 55087059 - 55223522 | | | | P |
| GSC-09 | RB1 | CCDS31973.1 | c.619C>T | p.Q207* | chr13:48934164 | C | T | 0.34 | P |
| | NF1 | CCDS42292.1 | c.1466A>G | p.Y489C | chr17:29541542 | A | G | 0.25 | P |
| | TERT | CCDS3861.2 | overlaps 1-kb region upstream | Promoter region | chr5:1295250 | G | A | 0.34 | P |
| | APC | CCDS4107.1 | c.1831G>A | p.G611S | chr5:112170735 | G | A | 0.28 | VUS |
| | PTEN | CCDS31238.1 | c.1010T>C | p.F337S | chr10:89720859 | T | C | 0.4 | LP |
| GSC-10 | TP53 | NM_000546 | c.772G>A | p.E258K | chr17:7577509 | C | T | 0.95 | P |
| | PIK3R1 | CCDS3993.1 | c.1042C>T | p.R348* | chr5:67588951 | C | T | 0.64 | P |
| | RB1 | CCDS31973.1 | c.2107-1G>A | Splice site | chr13:49037866 | G | A | 0.87 | P |
| | CHEK2 | CCDS33629.1 | c.599T>C | p.I200T | chr22:29121087 | A | G | 0.1 | VUS |
| | TERT | CCDS3861.2 | overlaps 1-kb region upstream | Promoter region | chr5:1295228 | G | A | 0.76 | P |
| | SPEN | CCDS164.1 | c.266G>A | p.R89Q | chr1:16199493 | G | A | 0.49 | VUS |
| | FAT4 | CCDS3732.3 | c.9973C>T | p.R3325C | chr4:126372144 | C | T | 0.52 | VUS |
| | PTEN | CCDS31238.1 | c.634+2T>A | Splice site | chr10:89712018 | T | A | 0.8 | LP |
| GSC-12 | TP53 | NM_000546 | c.586C>T | p.R196* | chr17:7578263 | G | A | 0.7 | P |
| | AMER1 | CCDS14377.2 | c.1921C>T | p.R641* | chrX:63411246 | G | A | 0.04 | P |
| | FANCA | CCDS32515.1 | c.2T>G | p.M1R | chr16:89883022 | A | C | 0.58 | P |
| GSC-13 | TP53 | NM_000546 | c.844C>T | p.R282W | chr17:7577094 | G | A | 0.49 | P |
| | TERT | CCDS3861.2 | overlaps 1-kb region upstream | Promoter region | chr5:1295250 | G | A | 0.53 | P |
| | RB1 | CCDS31973.1 | c.265-2A>C | Splice site | chr13:48916733 | A | C | 0.48 | LP |
| GSC-14 | TP53 | NM_000546 | c.733G>T | p.G245C | chr17:7577548 | C | A | 0.62 | P |
| | TP53 | NM_000546 | c.998G>A | p.R333H | chr17:7574029 | C | T | 0.2 | VUS |
| | RB1 | CCDS31973.1 | c.2211+1G>A | Splice site | chr13:49037972 | G | A | 0.57 | LP |
| | TERT | CCDS3861.2 | overlaps 1-kb region upstream | Promoter region | chr5:1295228 | G | A | 0.43 | P |
| | PTEN | CCDS31238.1 | c.1113delC | p.D371fs*45 | chr10:89725129 | AC | A | 0.6 | P |
| | FANCA | CCDS32515.1 | c.468G>C | p.L156F | chr16:89877169 | C | G | 0.3 | VUS |
| GSC-15 | TP53 | NM_000546 | c.880G>T | p.E294* | chr17:7577058 | C | A | 0.71 | P |
| | TERT | CCDS3861.2 | overlaps 1-kb region upstream | Promoter region | chr5:1295228 | G | A | 0.44 | P |
| | PIK3CA | CCDS43171.1 | c.2869A>C | p.T957P | chr3:178948097 | A | C | 0.44 | VUS |

| | | | | | | | | | |
|---|---|---|---|---|---|---|---|---|---|
| | PIK3R1 | CCDS3993.1 | c.454A>G | p.T152A | chr5:67569793 | A | G | 0.5 | VUS |
| | PALB2 | CCDS32406.1 | c.2803G>A | p.A935T | chr16:23635361 | C | T | 0.37 | VUS |
| GSC-16 | TP53 | NM_000546 | c.455C>T | p.P152L | chr17:7578475 | G | A | 0.26 | P |
| | TP53 | NM_000546 | c.686_687del | p.C229fs*10 | chr17:7577593 | TAC | T | 0.05 | P |
| | TERT | CCDS3861.2 | overlaps 1-kb region upstream | Promoter region | chr5:1295228 | G | A | 0.73 | P |
| | TP53 | NM_000546 | c.754_762del | p.L252_I254del | chr17:7577518 | TGATGGT GAG | T | 0.41 | P |
| | NF1 | CCDS42292.1 | c.6590_6591del | p.F2197fs*44 | chr17:29664545 | CTT | C | 0.32 | P |
| | PIK3R1 | CCDS3993.1 | c.1745+1G>- | Splice site | chr5:67591152 | TG | T | 0.64 | LP |
| | PTEN | CCDS31238.1 | c.95T>A | p.I32N | chr10:89653797 | T | A | 0.57 | LP |
| | RB1 | CCDS31973.1 | c.708delA | p.E237fs*27 | chr13:48934250 | CA | C | 0.61 | P |
| | BRCA1 | CCDS11453.1 | c.4987-2A>G | Splice site | chr17:41219714 | T | C | 0.29 | P |
| GSC-17 | TERT | CCDS3861.2 | overlaps 1-kb region upstream | Promoter region | chr5:1295228 | G | A | 0.46 | P |
| | EGFR | CCDS5514.1 | c.866C>A | p.A289D | chr7:55221822 | C | A | 0.84 | P |
| | PTEN | CCDS31238.1 | c.254-1G>A | Splice site | chr10:89692769 | G | A | 0.75 | LP |
| | JAK2 | CCDS6457.1 | c.842G>T | p.G281V | chr9:5054790 | G | T | 0.42 | VUS |
| GSC-18 | TERT | CCDS3861.2 | overlaps 1-kb region upstream | Promoter region | chr5:1295228 | G | A | 0.35 | P |
| | PTEN | CCDS31238.1 | c.808dupA | p.M270fs*28 | chr10:89720653 | C | CA | 0.26 | P |
| | NF1 | CCDS42292.1 | c.3655G>A | p.G1219R | chr17:29560178 | G | A | 0.43 | LP |
| | STAG2 | CCDS43990.1 | c.1734C>G | p.Y578* | chrX:123196968 | C | G | 0.44 | LP |
| | PLCB4 | CCDS54447.1 | c.1981C>A | p.L661M | chr20:9389810 | C | A | 0.6 | VUS |
| GSC-19 | TP53 | NM_000546 | c.839G>T | p.R280I | chr17:7577099 | C | A | 0.5 | P |
| | TERT | CCDS3861.2 | overlaps 1-kb region upstream | Promoter region | chr5:1295228 | G | A | 0.39 | P |
| | PTEN | CCDS31238.1 | c.358dupG | p.A120fs*6 | chr10:89692873 | T | TG | 0.5 | P |
| GSC-20 | TERT | CCDS3861.2 | overlaps 1-kb region upstream | Promoter region | chr5:1295228 | G | A | 0.49 | P |
| | NF1 | CCDS42292.1 | c.7549C>T | p.R2517* | chr17:29679366 | C | T | 0.3 | P |
| GSC-21 | TERT | CCDS3861.2 | overlaps 1-kb region upstream | Promoter region | chr5:1295228 | G | A | 0.29 | P |
| | PIK3R1 | CCDS3993.1 | c.1126G>C | p.G376R | chr5:67589138 | G | C | 0.21 | LP |
| | CHEK1 | CCDS8459.1 | c.1108T>G | p.W370G | chr11:12551441 3 | T | G | 0.52 | VUS |
| GSC-24 | TP53 | NM_000546 | c.743G>A | p.R248Q | chr17:7577538 | C | T | 0.5 | P |
| | PTEN | CCDS31238.1 | c.209+5G>A | Splice site | chr10:89685319 | G | A | 0.41 | LP |

| | | | | | | | | | |
|---|---|---|---|---|---|---|---|---|---|
| | TERT | CCDS3861.2 | overlaps 1-kb region upstream | Promoter region | chr5:1295228 | G | A | 0.39 | P |
| | SETD2 | CCDS2749.2 | c.4023G>A | p.W1341* | chr3:47162103 | C | T | 0.33 | LP |
| | RB1 | CCDS31973.1 | c.454_458del | p.L152fs*3 | chr13:48919288 | GTTGAA | G | 0.49 | LP |
| | NF1 | CCDS42292.1 | c.7189+2T>C | Splice site | chr17:29670155 | T | C | 0.47 | LP |
| GSC-25 | BRAF | CCDS5863.1 | c.1799T>A | p.V600E | chr7:140453136 | A | T | 0.22 | P |
| | TERT | CCDS3861.2 | overlaps 1-kb region upstream | Promoter region | chr5:1295228 | G | A | 0.41 | P |
| | PTEN | CCDS31238.1 | c.634+2T>G | Splice site | chr10:89712018 | T | G | 0.35 | LP |
| GSC-26 | TERT | CCDS3861.2 | overlaps 1-kb region upstream | Promoter region | chr5:1295250 | G | A | 0.39 | P |
| | PTEN | CCDS31238.1 | c.923G>A | p.R308H | chr10:89720772 | G | A | 0.45 | VUS |
| | NF1 | CCDS42292.1 | c.5907_5908del | p.R1970fs*6 | chr17:29661948 | CAA | C | 0.32 | LP |
| | FAM175A | CCDS3605.2 | c.367G>A | p.E123K | chr4:84391465 | C | T | 0.03 | VUS |
| | PTEN | CCDS31238.1 | c.106G>C | p.G36R | chr10:89653808 | G | C | 0.51 | LP |
| | PTPRB | CCDS44943.1 | c.4280A>G | p.Y1427C | chr12:70954603 | T | C | 0.31 | VUS |
| | NF1 | CCDS42292.1 | c.5177delT | p.L1726fs*8 | chr17:29653178 | CT | C | 0.29 | LP |
| GSC-31 | TERT | CCDS3861.2 | overlaps 1-kb region upstream | Promoter region | chr5:1295228 | G | A | 0.49 | P |
| | PTEN | CCDS31238.1 | c.830_838delinsT | p.T277fs*18 | chr10:89720679 | CATTCTTCA | T | 0.81 | P |
| | NF1 | CCDS42292.1 | c.6704+1G>T | Splice site | chr17:29664899 | G | T | 0.88 | P |
| | MAD2L2 | CCDS134.1 | c.368_369dup | p.R124fs*6 | chr1:11736160 | G | GGA | 0.05 | VUS |
| | SMARCA4 | CCDS12253.1 | c.4087_4092dupGAGGAG | p.E1363_E1364dup | chr19:11145715 | T | TGAGGAG | 0.42 | VUS |
| GSC-37 | EGFR | CCDS5514.1 | c.866C>T | p.A289V | chr7:55221822 | C | T | 0.35 | P |
| | TERT | CCDS3861.2 | overlaps 1-kb region upstream | Promoter region | chr5:1295250 | G | A | 0.61 | P |
| | PIK3R1 | CCDS3993.1 | c.1702C>T | p.P568S | chr5:67591109 | C | T | 0.45 | LP |
| | TP53 | NM_000546 | c.658T>G | p.Y220D | chr17:7578191 | A | C | 0.74 | P |

Supplemental Table 1. NGS results in gliosarcoma cases. VAF = Variant Allele Frequency. Interpretation represents pathogenic (P), likely pathogenic (LP) and variants of uncertain significance (VUS).
